# Shedding light (… electrons) on human bone ultrastructure with correlative on-axis electron tomography and energy dispersive X-ray spectroscopy tomography

**DOI:** 10.1101/2023.04.20.537681

**Authors:** Chiara Micheletti, Furqan A. Shah, Anders Palmquist, Kathryn Grandfield

**Author notes:** Corresponding authors: Prof. Anders Palmquist, Department of Biomaterials, Sahlgrenska Academy, University of Gothenburg, Gothenburg, Sweden, Prof. Kathryn Grandfield, Department of Materials Science and Engineering, McMaster University, Hamilton, ON, Canada.

## Abstract

Mineralized collagen fibrils are the building block units of bone at the nanoscale. While it is known that collagen fibrils are mineralized both inside their gap zones (intra-fibrillar mineralization) and on their outer surfaces (extra-fibrillar mineralization), a clear visualization of this architecture in three dimensions (3D), combining structural and compositional information over large volumes, but without compromising the resolution, remains challenging. In this study, we demonstrate the use of on-axis Z-contrast electron tomography (ET) with correlative energy-dispersive X-ray spectroscopy (EDX) tomography to examine rod-shaped samples with a diameter up to 700 nm prepared from individual osteonal lamellae in the human femur. Our work mainly focuses on two aspects: i) low-contrast nanosized circular spaces (“holes”) observed in sections of bone oriented perpendicular to the long axis of a long bone; and ii) extra-fibrillar mineral, especially in terms of morphology and spatial relationship with respect to intra-fibrillar mineral and collagen fibrils. From our analyses, it emerges quite clearly that most “holes” are cross-sectional views of collagen fibrils. While this had been postulated before, our 3D reconstructions and reslicing along meaningful two-dimensional (2D) cross-sections provide a direct visual confirmation. Extra-fibrillar mineral appears composed of thin plates that are interconnected and span over several collagen fibrils, confirming that mineralization is cross-fibrillar, at least for the extra-fibrillar phase. EDX tomography shows mineral signature (Ca and P) within the gap zones, but the signal appears weaker than that associated to the extra-fibrillar mineral, pointing towards the existence of dissimilarities between the two types of mineralization.

## 1. INTRODUCTION

At the nanoscale, bone is a composite whose two main components are type I collagen, a protein, and carbonated hydroxyapatite, a calcium-phosphate mineral [1]. Collagen molecules are quarter-staggered, creating overlap and gap zones which produce the characteristic 67 nm-periodic banding pattern seen in images of collagen fibrils [2,3]. In relation to collagen fibrils, mineralization is both intra-fibrillar, mainly in the gap zone [4], and extra-fibrillar (sometimes also referred to as inter-fibrillar), i.e., in the space outside or between the fibrils [5]. The presence of intra-fibrillar mineral in the gap zone is quite accepted, while extra-fibrillar mineral has received somewhat less attention, despite accounting for a significant fraction, if not even the majority, of the mineral phase according to some studies [6–11]. It has been proposed that the extra-fibrillar mineral is organized in stacks of four or more polycrystalline plates up to 200 nm in length, termed “mineral lamellae” [8–10,12]. Other studies identify mineralization as neither exclusively intra-or extra-fibrillar, but “cross-fibrillar”, encompassing multiple collagen fibrils [13], and indicate that the morphology of mineral is acicular, forming a needle-like habit, and stacking into larger aggregates [14]. In general, neither the spatial and temporal relationship between intra- and extra-fibrillar, nor their exact organization with respect to each other and to the collagen fibrils are fully understood.

One factor that complicates this is the hierarchical and varying organization of bone based on anatomical location. At the nanoscale, different patterns in bone ultrastructure have been described based on the orientation of mineralized collagen fibrils with respect to the image plane, stemming from their organization within the tissue as a whole. In longitudinal sections, i.e., sections cut parallel to the collagen fibril long axis, the characteristic banding pattern of collagen is noted, confirming that collagen fibrils lie within the image plane [8,13,15]. This pattern has been termed as longitudinal or “filamentous motif” [13], given the presence of mineral structures, identified as mineral lamellae by Schwarcz [12], which are elongated along the fibril length. In transverse sections, i.e., sections cut perpendicular to the collagen fibril long axis, seemingly empty or low-contrast circular to elliptical regions some tens of nanometers in diameter are observed dispersed between high-contrast mineral that curves or wraps around them [8,13,15,16]. This overall pattern has been given several names: as it resembles lace [8], the term “lacy motif” was adopted in [13], while others have referred to this view by its dominant rosette-like features [15].

It is now fairly well accepted that the filamentous and lacy motifs are mutually orthogonal two-dimensional (2D) projections of the same three-dimensional (3D) arrays of mineralized collagen fibrils. Evidence of this stems not just from images of sections oriented parallel and perpendicular to the long axis of a long bone where collagen fibril orientation is mostly known [8,15], but also from images collected at ±45° tilt angles using sections cut at 45° to the long axis of bone [8], and from electron tomography [9,13,15]. However, the content of the low-contrast circular regions is still debated. Is it collagen fibrils cut in cross-section, or do these circles represent nano-porosity, possibly occupied by non-collagenous matter, within bone? Cressey et al. first noted these low-contrast nanosized spaces surrounded by mineral in fossil human bone and modern sheep bone, and referred to them as “pores”, postulating that they are cross-sections of collagen fibrils [16]. Jantou et al., reinforced this theory when observing similar features in perpendicular sections of ivory dentin from elephant tusk [17]. This view was then adopted by more researchers in the bone field [8–10,12,15]. Grandfield et al. also noted nanosized circular features around and within mineral-rich rosettes, later to be interpreted in 3D as mineral ellipsoids [18], also suggesting that they correspond to collagen fibrils in cross-section [15]. Reznikov et al. disputed this interpretation, observing that the “lens-shaped”, “electron-transparent voids” typical of the lacy pattern do not match collagen fibrils in either size or distribution [13], while others proposed these are “unmineralized spaces” located within the collagen fibrils themselves and likely contain macromolecules [19]. While the studies cited so far are dominated by the use of transmission electron microscopy (TEM) and scanning TEM (STEM), similar structures have also been described by focused ion beam-scanning electron microscopy (FIB-SEM) tomography in the form of “nanochannels”, which are postulated to be involved in ion transport [20– 22]. Nanochannels were found to be most abundant at the periphery of circular mineral-rich areas that have a columnar shape in 3D [21], which most likely represent the same features termed mineral ellipsoids by other authors [18,23].

One of the challenges with identifying the contents of these spaces, as well as the organization and morphology of the mineral phase, in the studies to date is the limitations of electron microscopy imaging techniques. TEM has been the primary tool to investigate bone ultrastructure with nanoscale resolution [24]. While powerful, this tool is constrained with respect to the volume that can be analyzed, as TEM samples need to be electron transparent, usually <100 to 200 nm-thick. Moreover, S/TEM images are 2D projections, hence features along the sample thickness result overlapped in the final image. This limitation can be overcome by electron tomography (ET), where S/TEM images are collected over a range of tilt angles, and reconstructed by special algorithms to produce a 3D representation of the sample [25]. However, due to the holder geometry and spatial constraints within the sample chamber, the tilt range is typically restricted to ±70°, originating artifacts in the reconstruction due to a wedge-shaped unsampled region, the so-called “missing wedge” [25]. A more advanced type of ET, termed on-axis ET, uses specialized holders and rod-shaped samples to tilt over a ±90° range, or even allow for full 360° rotation, removing reconstruction artifacts due to the missing wedge in conventional ET [26,27]. Previous work on a bone-implant interface has demonstrated that on-axis ET can be effectively applied to the study of bone interfaces, leading to reconstructions with superior fidelity when high tilt ranges are used [28]. On-axis ET has also been combined with electron energy loss spectroscopy (EELS) tomography to correlate structural and compositional information, focusing on the chemical gradients at the bone-implant interface [29]. However, in both these studies [28,29], detailed analysis of bone nanoscale architecture was hindered by the more inconsistent orientation of collagen fibrils in bone formed around an implant placed in the maxilla, where collagen fibril organization reflects the complex loading patterns. On the other hand, distinct organizational motifs can be typically recognized in long bones such as the femur [8,13,15,30] and would be a more suitable candidate to investigate bone ultrastructure. To our knowledge, no work has combined ET with corresponding 3D analytical techniques, such as EELS or energy-dispersive X-ray (EDX) tomography, to simultaneously probe bone ultrastructure and composition.

As bone contains a significant amount of crystalline material in its mineral phase, ET is typically acquired in STEM mode to fulfill the so-called projection requirement [31]. This tomography mode is also known as *Z*-contrast tomography, as the contrast in images acquired with high-angle annular dark-field (HAADF) detectors in STEM strongly depends on composition, hence on the atomic number (Z). While ET provides 3D information with nanoscale resolution, the volume analyzed is still limited. Other electron microscopy techniques such as FIB-SEM can indeed examine larger volumes of material via a sequential slice and imaging protocol, but the trade-off is a loss in spatial resolution due to the probe size and a minimum slice thickness, perhaps at best 5-10 nm [32].

Herein, we combine on-axis Z-contrast ET of rod-shaped human bone samples, prepared from osteonal lamellae in the femoral cortex, with EDX tomography to gain more insights on the nature of the low-contrast regions characteristic of the lacy motif. By removing missing wedge artifacts and correlating spatial and compositional information with high resolution, we expose the contents of these regions. Hereinafter, we will refer to them as “holes” as in [8], leaving the quotation mark to reflect their low-contrast appearance and the debate on whether they are vacant or not. We also examine intra-fibrillar mineralization, as well as extra-fibrillar mineral morphology and its spatial organization with respect to collagen fibrils. This work applies a new methodology to correlate structural and compositional information to examine healthy human bone ultrastructure, which could be further expanded to the study of pathologies that alter bone nanoscale architecture.

## 2. MATERIALS AND METHODS

### 2.1. Sample preparation

#### 2.1.1. Bone sample

A sample of human femur from a 68-year-old male, without any known bone disease, obtained freshly-frozen, was fixed upon thawing in a solution of 4% glutaraldehyde in a 0.1 M cacodylate buffer for 7 days, and cut to obtain a transverse (i.e., perpendicular to the long axis of the femur) section of cortical bone using a slow speed diamond saw (IsoMet™, Buehler, IL, USA) under hydrated conditions (Figure S1A). The section was dehydrated in ethanol (70%, 80%, 90%, 95%, 100%) and embedded in Embed812 resin (Electron Microscopy Sciences, PA, USA). A detailed embedding protocol for mineralized bone samples is provided in [18]. The sample was polished (400, 800, 1200, and 2400 grit emery paper, followed by polishing cloth with a 50 nm diamond suspension), mounted on an Al stub with C and Ag tape, and coated with C (∼10-20 nm thickness). The sample was obtained with institutional ethical approval (HIREB No. 12– 085-T, McMaster University, ON, Canada).

#### 2.1.2. Site selection

Regions of interest (ROIs) for preparation of electron transparent samples for STEM imaging and ET were selected based on backscattered electron (BSE) images acquired in a SEM instrument (JEOL 6610LV, JEOL, MA, USA) operated at 10-15 kV (Figure S1B-C). ROIs for the rod-shaped samples were selected within a single osteonal lamella. These samples are numbered as *i* and *ii* hereinafter. ROIs for wedge-shaped samples were selected in three different orientations orthogonal to each other: sample *iii*, parallel to the osteonal axis and along an individual osteonal lamella; sample *iv*, parallel to the osteonal axis and across an osteonal lamella, i.e., centered on an individual lamella but crossing two inter-lamellar boundaries; sample *v*, perpendicular to the osteonal axis. Each sample was prepared in a different osteon, except samples *ii* and *v*. For a better understanding of the ROIs locations, please refer to the schematic in Figure S1D-K.

#### 2.1.3. Preparation of rod-shaped samples for ET

Two rod-shaped samples for ET experiments (samples *i* and *ii*) were prepared by *in situ* lift-out followed by annular milling in a dual-beam FIB instrument (Zeiss NVision 40 and Zeiss Crossbeam 350, Carl Zeiss AG, Germany) equipped with a 30 kV Ga^+^ ion column following published protocols [33]. Briefly, after depositing a layer of W over a previously selected 3 × 3 μm^2^ ROI, trenches were milled around the ROI at 30 kV with currents in the 6.5-65 nA range (lower values when approaching the ROI). The sample was attached to the micromanipulator, lifted-out and attached to a 1 mm cartridge for an on-axis rotation tomography holder (Model 2050, Fischione Instruments, PA, USA). The sample was milled with annular milling patterns at 30 kV and progressively lower currents (80-100 pA, 40-50 pA).

#### 2.1.4. Preparation of wedge-shaped samples

Wedge-shaped samples for STEM experiments (samples *iii-v*) were prepared in a dual-beam FIB instrument (Zeiss NVision 40, Carl Zeiss AG, Germany) equipped with a 30 kV Ga^+^ ion column.

##### Samples parallel to the osteonal axis

Samples *iii* and *iv* were prepared by *in situ* lift-out following published protocols [34]. Briefly, after depositing a layer of W over a previously selected 12 × 2 μm^2^ ROI, trenches were milled around the ROI at 30 kV with currents in the 6.5-45 nA range (lower values when approaching the ROI). The sample was attached to the micromanipulator, lifted-out and attached to a Cu grid. In each sample, three ∼2 μm-wide windows were thinned to electron transparency at 30 kV and progressively lower currents (150, 80, and 40 pA), leaving thicker supports in between. Final beam polishing was completed at 5 kV and 60 pA to limit Ga^+^ implantation and amorphization damage.

##### Sample perpendicular to the osteonal axis

To avoid re-mounting the bone sample, sample *v* was prepared following a *plan-view* lift-out protocol similar to [35]. A layer of W was deposited over a previously selected 15 × 15 μm^2^ ROI. Trenches were milled around the ROI at 30 kV with currents in the 6.5-45 nA range (lower values when approaching the ROI). The sample was attached to the micromanipulator, lifted-out, and attached to a Cu grid mounted horizontally in a specialized stub with two pins located at 90° from each other (this holder is described in [33]). The stub was removed from the holder, rotated by 90°, and repositioned in a standard holder so that the Cu grid was now oriented vertically, as in conventional *in situ* lift-outs. The sample was thinned to electron transparency as described above for the samples *iii* and *iv*.

### 2.2. STEM imaging and on-axis ET

STEM imaging and ET experiments of the rod-shaped samples *i* and ii were completed in a S/TEM instrument (Talos 200X, Thermo Fisher Scientific, MA, USA) equipped with four in-column silicon drift detectors (Super-X detector) and operated at 200 kV. The same instrument was used to acquire STEM images of wedge-shaped samples *iv* and *v*, while STEM images of sample *iii* were acquired in a Titan 80-300 LB (Thermo Fisher Scientific, MA, USA) operated at 200 kV. For on-axis ET, an on-axis rotation tomography holder (Model 2050, Fischione Instruments, PA, USA) was used. ET was completed using the STEM Tomography application in Velox (Thermo Fisher Scientific, MA, USA), acquiring multiple signals simultaneously (HAADF-STEM and BF-STEM, HAADF-STEM and EDX) with automated focusing and image shifting based on the HAADF-STEM image. In each rod-shaped sample, two tilt series were acquired in two different locations (marked by rectangles in Figure 1). In sample *i*, two tilt series were acquired over a ±90° range, completely removing missing wedge artifacts, while in sample *ii* the tilt range was restricted to a ±85° range to simplify acquisition and reconstruction. A summary of acquisition conditions is reported in Table 1. In all samples, EDX maps of Ca, P, C, and N were acquired over 10 frames with a 50 μm dwell time per pixel and smoothing to denoise (Gaussian blur with σ = 2; 5-7 pixel average). Background-subtracted maps were generated automatically in the Velox software and recorded at each tilt step.

**Table 1.**
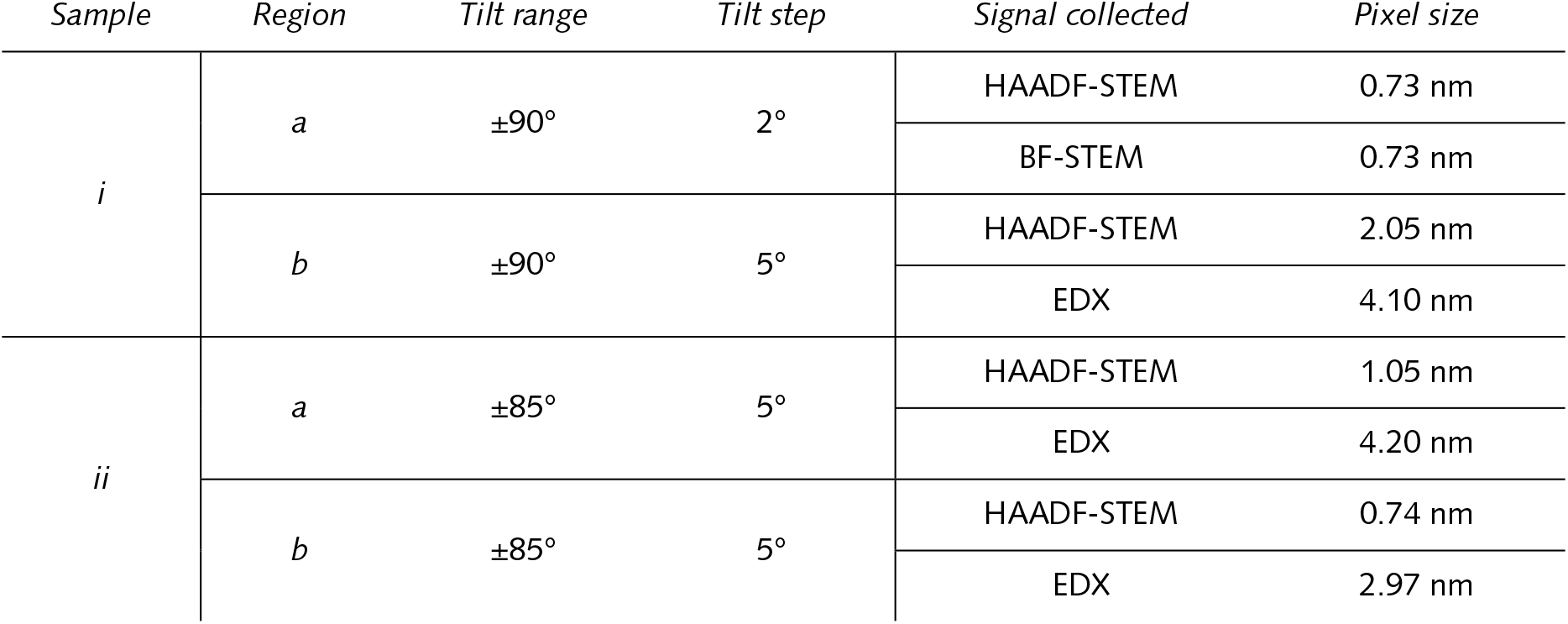
ET acquisition conditions for the regions examined in the rod-shaped samples *i* and *ii*.

| Sample | Region | Tilt range | Tilt step | Signal collected | Pixel size |
| --- | --- | --- | --- | --- | --- |
| *i* | *a* | $\pm 90^\circ$ | $2^\circ$ | HAADF-STEM | 0.73 nm |
|  |  |  |  | BF-STEM | 0.73 nm |
| | *b* | $\pm 90^\circ$ | $5^\circ$ | HAADF-STEM | 2.05 nm |
|  |  |  |  | EDX | 4.10 nm |
| *ii* | *a* | $\pm 85^\circ$ | $5^\circ$ | HAADF-STEM | 1.05 nm |
|  |  |  |  | EDX | 4.20 nm |
| | *b* | $\pm 85^\circ$ | $5^\circ$ | HAADF-STEM | 0.74 nm |
|  |  |  |  | EDX | 2.97 nm |

**Figure 1.**
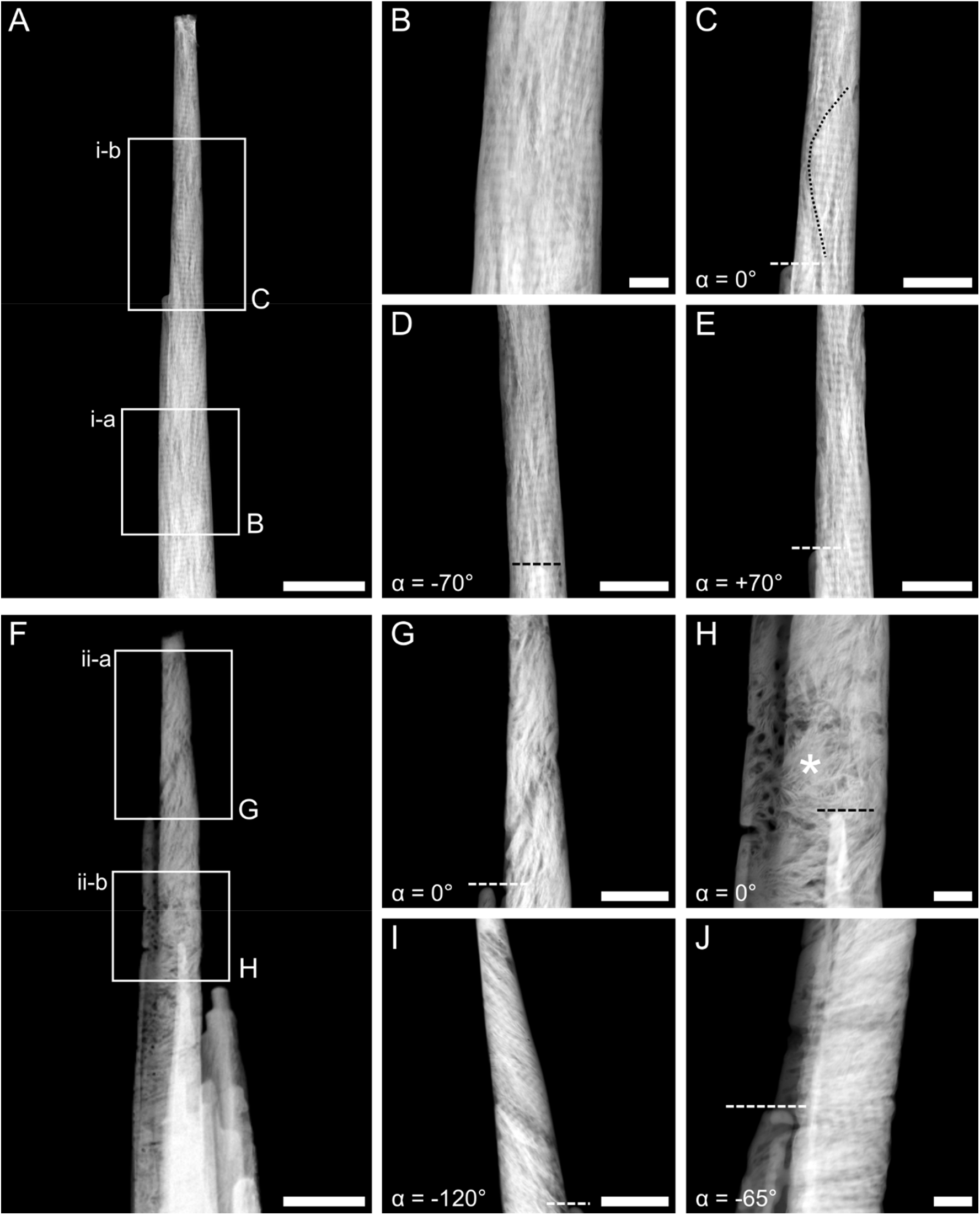
HAADF-STEM images of rod-shaped samples. A) Overview image of the rod-shaped sample *i*. B-C) Higher magnification images where the tilt series *i-a* (B) and *i-b* (C) were acquired. In sample *i*, collagen banding is visible throughout the length of the sample and at different tilt angles (α). An incomplete mineral ellipsoid can also be noted in C, where its boundary is marked by the black dotted line. D-E) Images acquired in the same region as C but tilted to -70° (D) and +70° (E). F) Overview image of the rod-shaped sample *ii*. G-H) Higher magnification images where the tilt series *ii-a* (G) and *ii-b* (H) were acquired. In sample *ii*, collagen is present both out-of-plane (G-H) and in-plane (I-J) based on the tilt angle. A mineral-rich region resembling a rosette surrounded by circular dark features can be noted in H (marked by *). I) Collagen banding is more apparent when imaging the region in G at a tilt angle equal to -120°. J) Collagen banding is present instead of the rosette in H when imaging at a different tilt angle (−65°). The dotted lines in C-E and G-J mark the same feature seen at different tilt angles for ease of interpretation. Please note that, given the cylindrical geometry of the samples and the rotation holder used, the tilt angle α represents a relative and not absolute reference. Scale bars are 1 μm in A and F, 200 nm in B, H, and J, and 500 nm in C, D, E, G, and I.

### 2.3. Data reconstruction, visualization, and analysis

All the tilt series were aligned by cross-correlation, followed by manual adjustment of the tilt axis shift, and reconstructed using a simultaneous iterative reconstruction technique (SIRT) with 25 iterations in Inspect 3D (Thermo Fisher Scientific, MA, USA). BF-STEM and EDX tilt series were aligned based on the HAADF-STEM tilt series. The reconstructed electron tomograms were visualized and analyzed in Dragonfly (Object Research Systems, QC, Canada), as described below.

#### Analysis of the “holes”

“Holes” (dark in HAADF-STEM and bright in BF-STEM reconstructed slices) were segmented as follows: 1) a ROI containing both these features and the background (i.e., the region around the rod-shaped sample included in the field of view of the detector) was segmented using the “Define range” operation; 2) another ROI was created by inverting the ROI in point 1 (note: this ROI mostly corresponds to mineralized regions); 3) the ROI in point 2 was filled using the “Fill inner areas” operation (applied in 2D in x, y, and z); 4) a ROI was created by a Boolean intersection operation between the ROIs in points 1 and 3 to isolate the dark (in HAADF-STEM)/bright (in BF-STEM) features contained within the sample (i.e., the “holes”). This was necessary to separate the “holes” from the background. The ROI in point 4 was further refined using the “Process islands” operation to remove mislabeled voxels. The size of the “holes” was evaluated by applying the “Volume thickness map” operation to the ROI in point 4. The ROI in point 4 was split in separate ROIs using the “Connected components – Multi ROI (6-connected)” operation. Intensity profiles along specific features were extracted using the “Ruler” tool. The distance between local minima and maxima in the intensity profile was evaluated with the function “argrelextrema” in the “scipy.signal” library in Python 3.8.10. The average distance between peaks was evaluated as the arithmetic mean between the average distance between local minima and maxima.

#### Analysis of mineral

A “coarse” segmentation of the mineral phase based on grey-levels was performed using the “Define range” operation. This ROI was split in separate ROIs using the “Connected components – Multi ROI (6-connected)” operation. Some representative mineral structures (3-5 in each tomogram) were manually segmented in the reconstructed slices with the “Brush” tool with local Otsu thresholding, selecting the upper Otsu range only. In tomograms *i-a*/*b* and *ii-a*, mineral structures were assumed to be oriented with their length predominantly within the xy planes and their width in the yz planes (refer to Figure S1 for coordinate system convention). For these samples, variation in mineral length and thickness was evaluated with the “Slice analysis” module by computing the perimeter (2p) and thickness (t) of the segmented feature in each xy slice, and estimating the length (l) by subtracting the thickness from the semi-perimeter (l = p - t). The width was estimated by multiplying the number of xy slices where the feature was segmented by the voxel size in z. In tomogram *ii-b*, mineral structures were assumed to be oriented with their width predominantly within the xy planes and their length in the xz planes (refer to Figure S1 for coordinate system convention). For these samples, variation in mineral width and thickness was evaluated with the “Slice analysis” module by computing the perimeter (2p) and thickness (t) of the segmented feature in each xy slice, and estimating the width (w) by subtracting the thickness from the semi-perimeter (w = p - t). The length was estimated by multiplying the number of xy slices where the feature was segmented by the voxel size in z.

#### Analysis of EDX maps

EDX tomograms of Ca, P, C, and N were segmented to select the most intense signal only (for each element independently). A Boolean intersection operation was applied between the segmented EDX signal for each element and the segmented “holes” to quantify the content (expressed as number of voxels) of each element within the segmented “holes” only.

### 2.4. Simulation of ion beam-sample interaction

Implantation of Ga^+^ in bone during FIB sample preparation was simulated using the SRIM (Stopping and Range of Ions in Matter) software [36]. Bone was considered as a compound of 70 wt% mineral (hydroxyapatite) and 30 wt% organic (type I collagen) [37]. The mineral phase was approximated as made of 47.8 wt% Ca and 22.2 wt% P, considering a Ca/P atomic ratio of 1.67 for hydroxyapatite [38]. The organic phase was approximated as solely composed of C. Bone density was set equal to 1.8 g/cm^3^ [39].

### 2.5. Estimation of X-ray absorption

Absorption of X-rays during EDX tomography experiments was estimated considering the following attenuation law [40]:

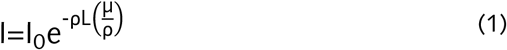

where I/I_0_ is the fraction of X-rays not absorbed, L is the path length, ρ is the density of the material, and the ratio μ/ρ represents the mass attenuation coefficient of the material at a certain energy [40]. A maximum path length (L) of 700 nm was considered, since this corresponds to the maximum diameter of the rod-shaped samples in the regions where EDX tomography was completed. Bone density (ρ) was set equal to 1.8 g/cm^3^ [39], as in the simulation of ion beam-sample interaction. The mass attenuation coefficient (μ/ρ) was considered to be equal to 3.781 × 10^3^ cm^2^/g, which is the tabulated value for cortical bone at 1 keV [41]. This is the tabulated value closest to that of the lowest X-ray energy for the elements we mapped with EDX tomography (C Kα = 0.277 keV).

## 3. RESULTS AND DISCUSSION

### 3.1. Collagen fibril orientation in rod-shaped samples

Two quasi-cylindrical, slightly conical samples (Figure 1) were obtained by FIB annular milling. The samples, referred to as *i* and *ii*, have a diameter ranging from approximately 200-300 nm at the top to 700 nm at the bottom, and a length that retained electron transparency of 8 μm and 4 μm for *i* and *ii*, respectively. Sample *i* presents collagen banding perpendicular to the long axis of the rod irrespective of the tilt angle, confirming collagen fibrils laying roughly in-plane at all tilts and co-aligned with the long axis of the rod (Figure 1A-E). The trace of some mineral ellipsoids is faintly distinguishable (Figure 1C, black dotted line), though a full ellipsoid was not captured. On the other hand, in sample *ii*, two different motifs are visible (Figure 1F-J). While a more disorganized region is distinguishable in the top half of the sample (Figure 1G), this disappears upon tilting and in-plane collagen fibrils can be noted instead (Figure 1I). An area with a mineral-rich rosette-like pattern dominates the bottom half of the sample with collagen fibrils appearing out-of-plane (Figure 1H) at some angles, then in-plane at other angles (Figure 1J). The banding patterns are nearly orthogonal to one another in the upper and lower regions, and this juncture likely represents that the boundary between two lamellae was crossed along the length of sample *ii*.

For comparison to conventional wedge-shaped TEM samples, sample *i* well corresponds to the longitudinal motif of in-plane collagen in sample *iii* (Figure S2A-B). Conversely, the more inconsistent and angle-dependent orientation of the collagen fibrils in sample *ii* resembles the patterns observed in samples *iv* and *v* (Figure S2C-F), with transitions from longitudinal to lacy motif, hence indicating variations in fibril organization within the osteonal lamella selected for sample *ii* preparation.

### 3.2. What are the “holes”?

#### Co-locating “holes” with collagen banding

Electron tomograms were acquired in the boxed regions shown in Figures 1B (tomogram *i-a*), C (tomogram *i-b*), G (tomogram *ii-a*), and H (tomogram *ii-b*). For tomograms *i*-*b* and *ii-a*/*b* we simultaneously acquired EDX maps for C and N as markers of the organic phase, and Ca and P as markers of the mineral phase.

Electron tomograms give accurate 3D volumes that can be resliced in any orientation. By reslicing, the reconstructions clearly confirmed the correspondence between the longitudinal and lacy motifs in mutually orthogonal planes. Specifically, in Figure 2, the banding pattern is present in the reconstructed slices in both xy and yz planes, while “holes” are visible in the xz plane instead, corresponding to similar 2D S/TEM images from either longitudinal or lacy motifs (Figure 2D,F). The reciprocity between “holes” and collagen fibrils is probed in more detail in Figure 3, where the banding pattern is noted in the reconstructed xy and yz slices in correspondence to a specific “hole” marked by a box in the xz plane (Figure 3B-E). The intensity profile extracted in the regions marked by the rectangles in Figures 3C and D confirms the presence of a periodic variation in grey-levels, with an average distance between the peaks in the profile of 66.5 nm and 58.5 nm in panels C and D, respectively (Figure 3F), quite close to the expected 67 nm for D-spacing in collagen fibrils. Another example of collagen banding pattern orthogonal to the “holes” is presented in Figure S3.

**Figure 2.**
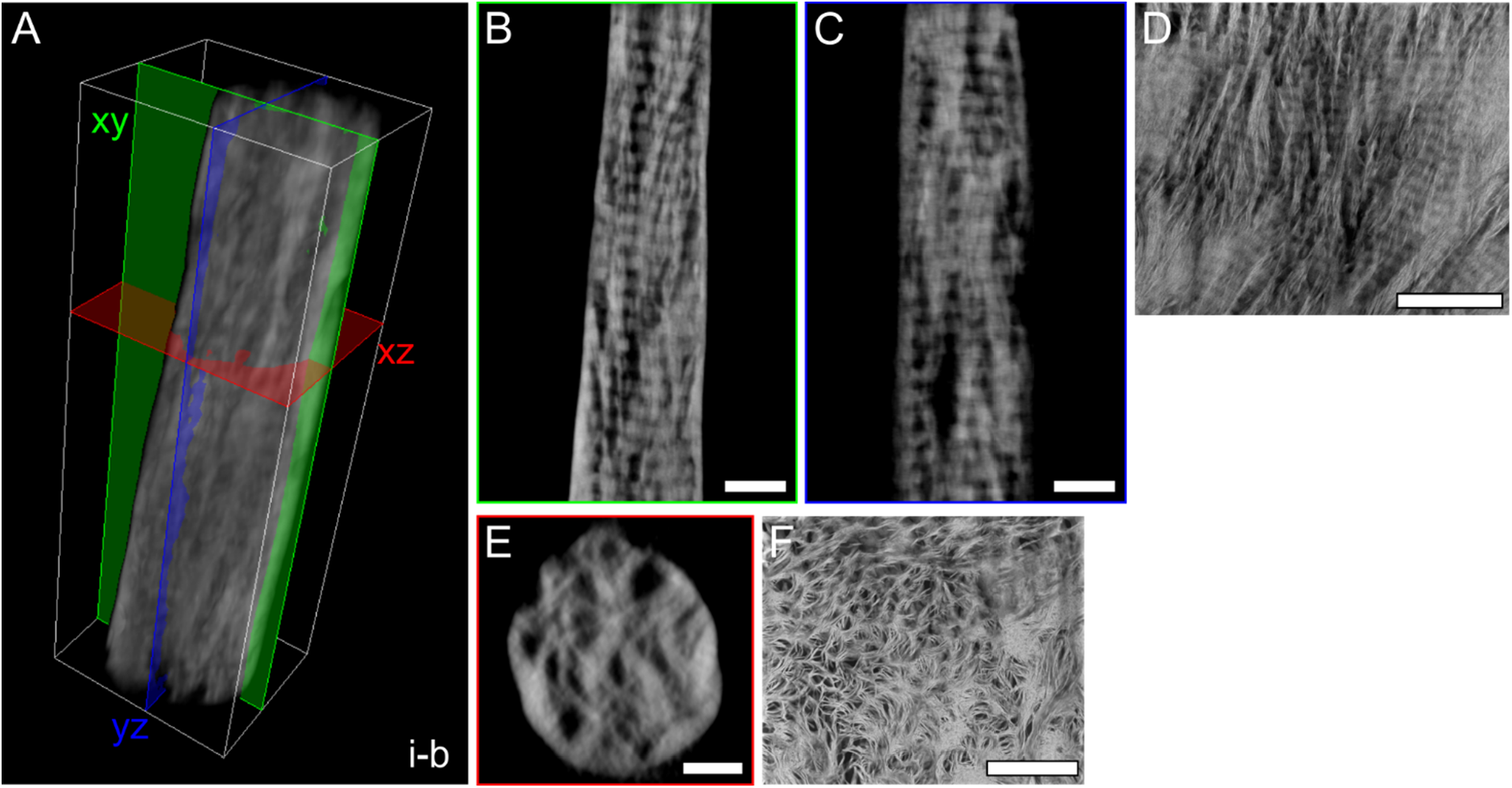
Banding pattern and “holes” are seen in mutually orthogonal views. A) 3D reconstruction of tomogram *i-b*, where three mutually orthogonal planes are marked (xy: green; yz: blue; xz: red). B) A representative reconstructed slice in the xy plane, corresponding to the green plane in A. C) A representative reconstructed slice in the yz plane, corresponding to the blue plane in A. D) Longitudinal motif shown in a HAADF-STEM image of the wedge-shaped sample *iii*, prepared parallel to the osteonal axis within a single lamella such that collagen fibrils are in the image plane. This motif is analogous to that of B and C. E) A representative reconstructed slice in the xz plane corresponding to the red plane in A, showing dark circular features or “holes*”*. F) HAADF-STEM image of the wedge-shaped sample *v* (prepared perpendicular to the osteonal axis), where the presence of “holes” and encircling mineral is analogous to the view in E. Scale bars are 200 nm in B, C, D, and F, and 100 nm in E. A scale bar is not provided in A as the 3D representation is not an orthographic projection; the dimensions (x, y, z) of the white box are 555.55 × 1611.30 × 457.15 nm^3^.

**Figure 3.**
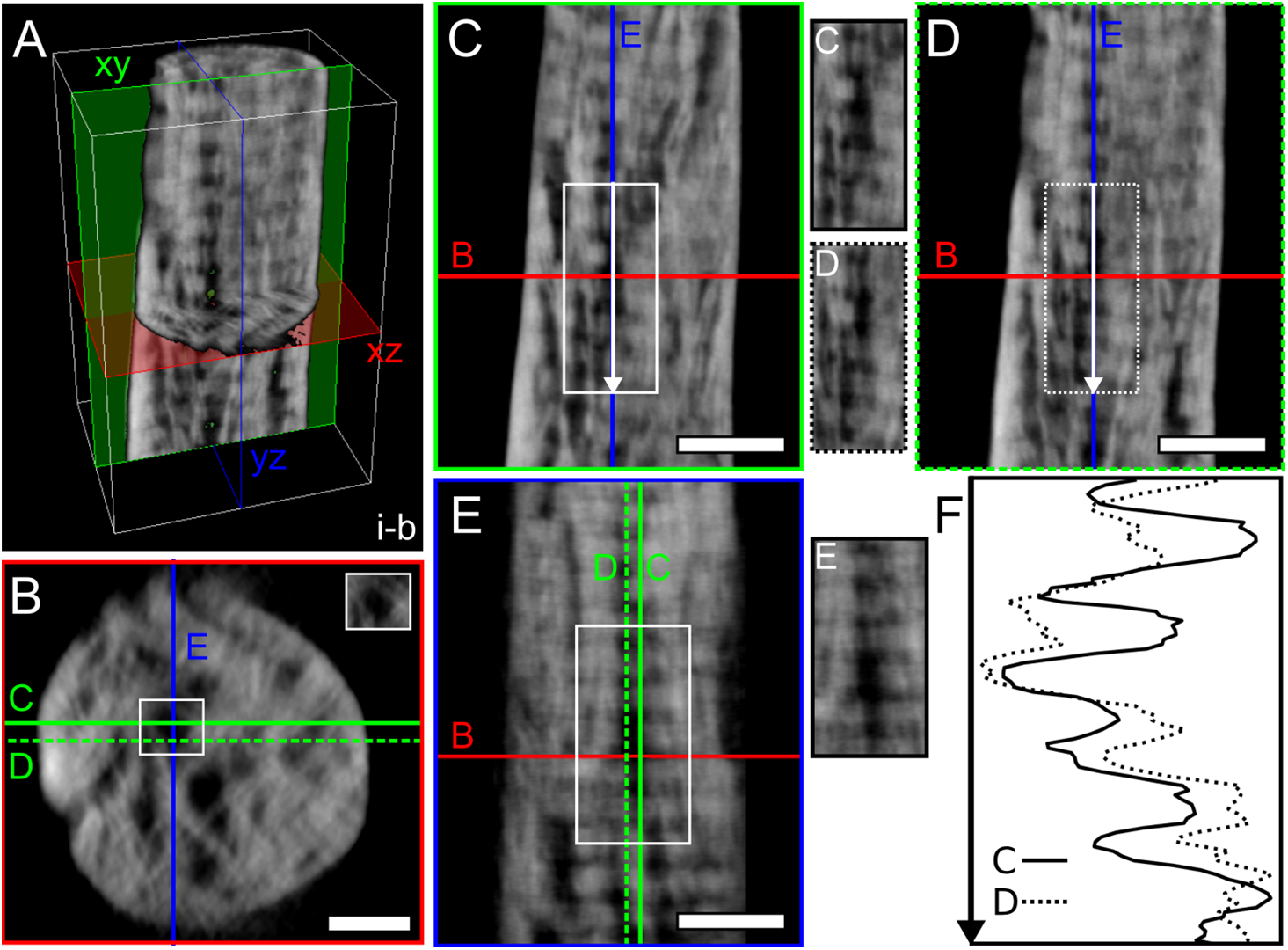
Co-localization of “holes” and banding pattern. A) 3D rendering of a section of tomogram *i-b*. B) A representative reconstructed slice in the xz plane where “holes” can be noted, originating a pattern similar to the lacy motif. The “hole” marked by the white rectangle was examined in the two planes orthogonal to xz (red), i.e., xy (green) and yz (blue). C) Slice in the xy plane corresponding to the solid green line in B. D) Slice in the xy plane corresponding to the dashed green line in B. The xy slices in C and D are 22.55 nm away from each other in the z direction. E) Slice in the yz plane corresponding to the blue line in B. F) The intensity profile extracted from 350 nm-long lines marked by arrows in C and D confirms the presence of collagen banding pattern (average distance between minima and maxima equal to 66.5 nm and 58.5 nm in C and D, respectively). An unmarked image of the regions marked by rectangles in C, D, and E is provided next to each panel. Scale bars are 100 nm in B, and 200 nm in C, D, and E. A scale bar is not provided in A as the 3D representation is not an orthographic projection; the dimensions (x, y, z) of the white box are 555.55 × 865.10 × 457.15 nm^3^.

The clear correspondence between banding pattern and “holes” in mutually orthogonal views enforces that the “holes” are indeed filled with collagen fibrils that are seen in cross-section. This is especially apparent in the tomograms acquired in sample *i* (Figure 1B-C), due to the optimal orientation of the banding pattern with respect the long axis of the sample. Yet, it follows that if collagen indeed fills the “holes”, the cross-sectional views should differ in a gap versus an overlap zone in a mineralized sample, assuming that at least some of the mineral is located in the gap zone (intra-fibrillar). To further demonstrate this, we extracted xz slices within gap and overlap zones from an area where the banding pattern is well resolved over several bands, especially when adjusting the grey-level range to better encompass low-contrast elements (Figure 4). When examining the cross-sections in the xz plane, it becomes evident that the darkest (blackest) “holes” are cross-sectional views of overlap zones, hence explaining the nature of their low-contrast appearance, while the gap zones have some matter and a brighter grey-level detected (Figure 4D).

**Figure 4.**
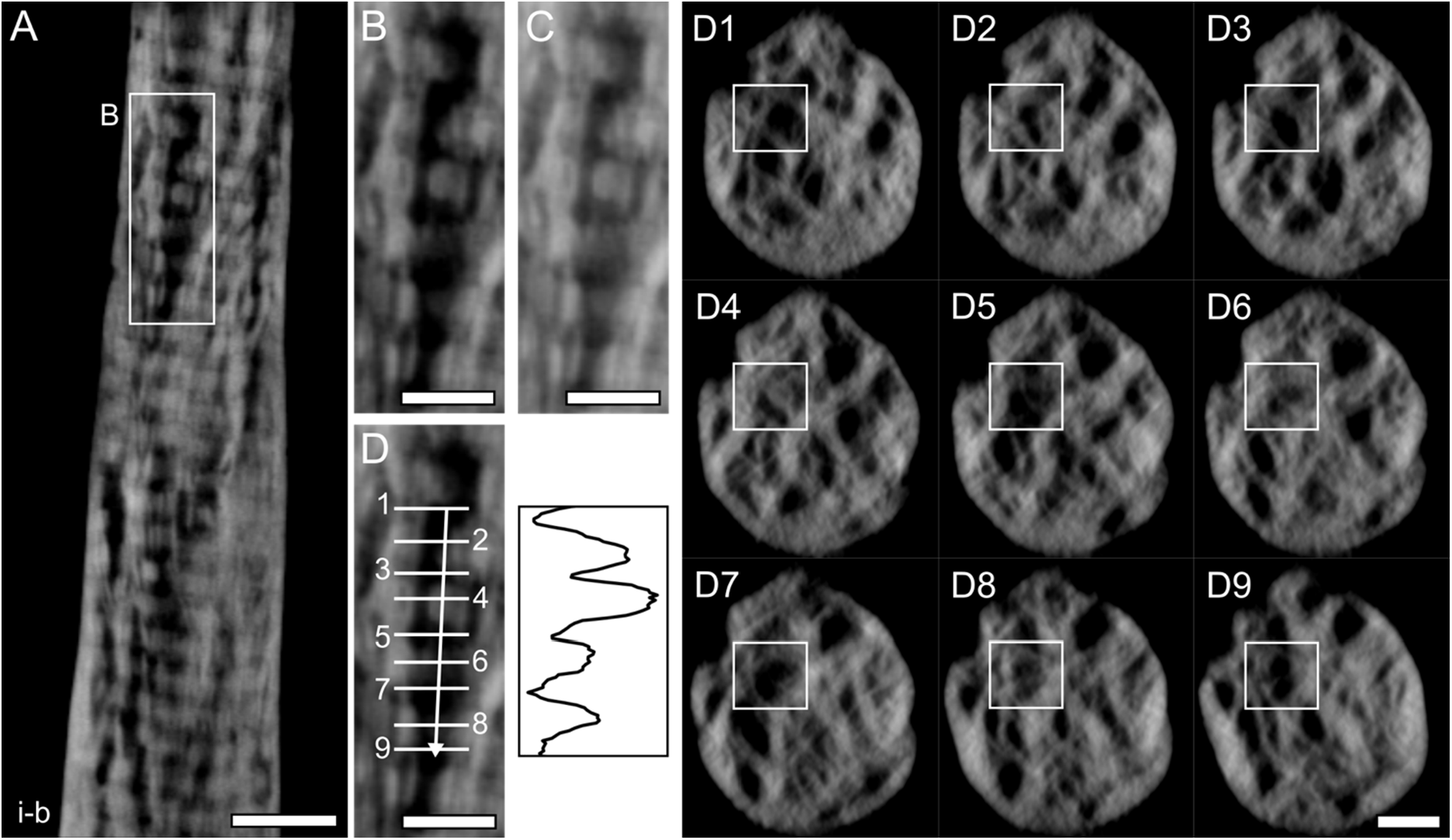
Dark “holes” are cross-sections of overlap zones. A) A representative reconstructed slice in the xy plane in tomogram *i-b*. B) Banding pattern can be noted, but is obscured by the low contrast in some regions. C) The banding pattern is more noticeable when grey-levels are adjusted to include lower values. D) Different cross-sections (xz plane) are examined in gap (even numbers) and overlap (odd numbers) zones in the xy plane. “Holes” are mostly visible when cross-sectioning along the overlap regions (the “hole” marked in D1-9 corresponds to the region shown in the xy plane in D). The histogram profile extracted along the white arrow in D confirms the presence of periodicity (61-66 nm). Scale bars are 200 nm in A, and 100 nm in B, C, and D.

In summary, as the banding pattern is the dominant motif in the xy and yz planes of our ET reconstructions, it seems natural to conclude that most “holes” in the lacy motif (xz plane) are cross-sections of collagen fibrils, confirming what was first proposed by Cressey et al. [16] and supported by other authors [8–10,12,15,17]. If this was not the case, more areas devoid of collagen fibrils, i.e., without any clear banding pattern and pure black, should be apparent in the xy/yz planes. Some of these areas were indeed noted (Figure S4A-C), but in a very limited amount, which could not then explain the extensive presence of “holes” in the xz view. In the locations where the banding pattern cannot be resolved, it cannot be truly determined whether collagen fibrils are indeed present but cross-sectioned along a plane that makes the banding less evident, or if they correspond to true nano-porosity or areas where non-collagenous elements are located. Recent work by Macías-Sánchez et al. in similar cross-sectional views of mineralizing turkey leg tendon proposed that unmineralized spaces exist within collagen fibrils and macromolecules are likely contained therein [19].

#### Why do “holes” appear empty in most S/TEM images?

In ET reconstructions, most “holes” in the xz plane correspond to the banding pattern in the xy plane. However, in previous works, “holes” seemed largely empty in S/TEM images of conventional wedge-shaped TEM samples. This vacant appearance of the “holes” in the lacy motif has been previously attributed to the preferential erosion of the organic phase by the ion beam during sample preparation [8]. In EELS experiments in ivory dentin, Jantou et al. observed that t/λ (relative thickness) was never equal to zero in the “holes”, indicating that they are not empty [42]. Furthermore, Lee et al. showed also by EELS that the “holes” were rich in C and N [43]. To confirm the hypothesis of ion erosion, we simulated the interaction between Ga^+^ ion and bone, which yielded an implantation depth up to nearly 50 nm at 30 kV. Hence, ion erosion justifies the vacant nature of “holes” in wedge-shaped samples that only have a 100-200 nm thickness, considering that thinning to electron transparency is typically performed on both sides of the sample. Conversely, the ion beam cannot erode the collagen fibrils in central regions of the thicker 300-700 nm rod-shaped samples shown here. Ion erosion could explain the lack of banding pattern in correspondence with some “holes” located near the outer surface of the rod-shaped samples presented herein (Figure S4A,D), but overall the larger thickness of the samples ensures that collagen was not eroded away, enabling its clear visualization in planes orthogonal to the “holes”. This confirms that nearly all “holes” seen in so-called lacy or rosette motifs of bone are indeed collagen fibrils in cross-section.

#### Size and organization of the “holes” in 3D

The size and arrangement of the “holes” were comprehensively analyzed in 3D after segmenting the tomograms based on grey-levels, which included those “holes” that appear rather low-contrast in xz slices. While “holes” display a circular/elliptical shape in S/TEM images of the lacy motif and in the reconstructed xz slices in tomograms *i-a*/*b* (and in well-oriented cross-sections in tomogram *ii-a*), their visualization in 3D reveals that they are elongated rod-like features mainly aligned perpendicular to the banding pattern. This is yet another similarity with and evidence of them being collagen fibrils. Therefore, segmentation of low-contrast “holes” likely captures fragments of mineralized collagen fibrils less masked by the brighter contrast of the mineral phase.

The size of the segmented “holes” was evaluated using the “Volume thickness map” operation in the Dragonfly software, yielding an average diameter of 22.9 ± 4.6 nm over the four tomograms (Figure 5A, Figure S5, Table S1). Specifically, the average diameter is in the 22.9-26.4 nm range in tomograms *i-b* and *ii-a/b*, but is reduced to 16.4 nm in tomogram *i-a*. At a closer inspection, it appears that a higher number of smaller features below 10 nm are captured in the segmented “holes” in tomogram *i-a* (Figure S5). Overall, the size of the segmented “holes” does not correspond to what is commonly indicated for mineralized collagen fibrils, i.e., 80-120 nm [1,44,45]. On the other hand, “holes” in out-of-plane S/TEM images of bone have also been described as 20-50 nm in size [8,12,13,15]. Mineralized collagen fibrils with diameters as small as 30 nm have been reported using atom probe tomography [46]. Sample preparation artifacts such as shrinkage (up to 27% [47]) or fibrils cross-sectioned along planes not perfectly perpendicular to their long axis could also explain our smaller measurements. Hence, considering the co-localization of “holes” and banding pattern in mutually orthogonal slices, it appears inaccurate to rule out that “holes” are cross-sections of collagen fibrils solely based on size considerations.

**Figure 5.**
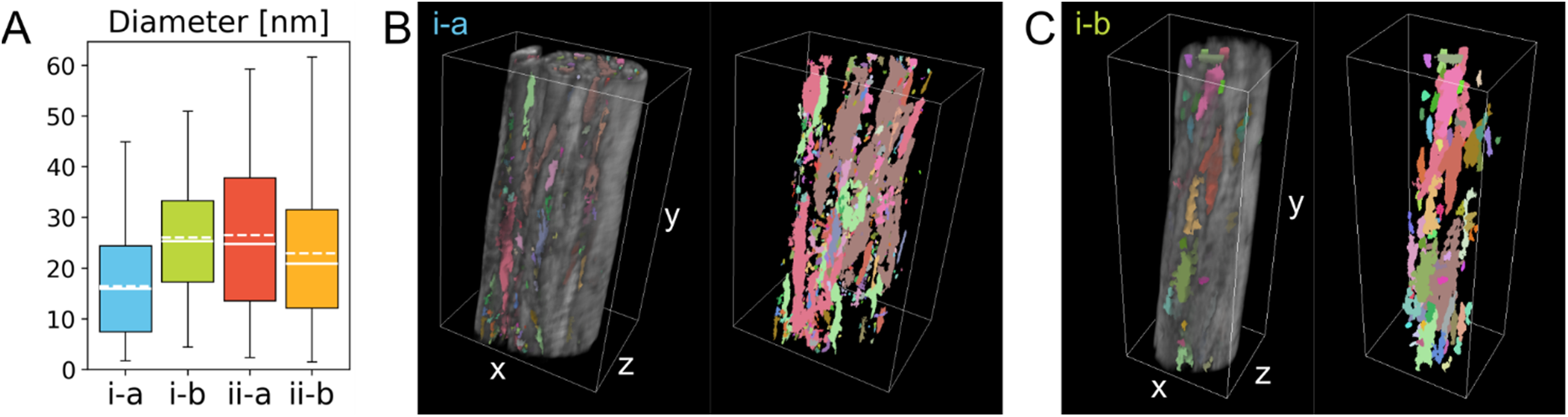
Size and connectivity of the “holes”. A) Box plot of the diameter of the segmented “holes” in each tomogram. Each box represents 25-75% of the data, while the capped lines indicate the entire range of the data. Median and mean values are marked by solid and dashed lines, respectively. B) 3D rendering of tomogram *i-a* with the segmented “holes”, shown in different colours to represent their disconnected nature. C) Similar visualization as B for tomogram *i-b*. A scale bar is not provided in B and C as the 3D representation is not an orthographic projection; the dimensions (x, y, z) of the white box are 879.65 × 1416.20 × 620.50 nm^3^ in B, and 555.55 × 1611.30 × 457.15 nm^3^ in C.

Interestingly, the average diameter of “holes” in our segmentations is well within the 10-50 nm range reported for nanochannels [20– 22]. However, some discrepancies also exist. Most nanochannels in bone have a certain degree of interconnectivity (3-4 nodes) [21] and are micrometres in length [22]. On the other hand, our connectivity analysis of the segmented “holes” using the “Connected components – Multi-ROI” operation in the Dragonfly software, which splits the segmented ROI in multiple ROIs based on their degree of connection, revealed that “holes” are mostly disconnected from each other (Figure 5B-C, Figure S6). Additionally, no segmented “holes” extending over the entire sample length are present. Conversely, most appear rather short, with a length of at most ∼500 nm in the longest continuous segments. Overall, more evidence indicates that “holes” are collagen fibrils, rather than other hole-like features like nanochannels. However, it is possible that some regions we noted where the banding pattern is not present do indeed correspond to nanochannels.

In the region of tomogram *i-b* where a partial mineral ellipsoid can be seen (Figure 1C in HAADF-STEM), not many segmented “holes” are present within the ellipsoid, but they seem most abundant around its periphery (Figure S7). This is probably due to the mineral-rich nature of the ellipsoids, especially in their core [30], which obscures low-contrast elements, resulting in fewer dark “holes” being detected within. Similarly, “holes” (already interpreted as collagen fibrils in that study) were mostly noted on the periphery of rosettes in 2D HAADF-STEM images [15]. On the other hand, the lower mineral content in between ellipsoids, likely due to the accumulation of mineral inhibitors [48], makes the “holes” (i.e., the collagen fibrils) more noticeable in the images due to contrast effects. By analogy, since both nanochannels [21] and unmineralized spaces [19] are noted at the periphery of ellipsoids (called spherulites in [19]), it is indeed possible that at least some of these features do indeed correspond to cross-sections of collagen fibrils.

#### Correlative 3D compositional information with EDX tomography

We completed correlative HAADF-STEM and EDX tomography experiments to confirm whether “holes” are rich in organic components, namely C and N. Unfortunately, the spatial distribution of C and N does not provide resolutive information in this regard, as the maps appear quite noisy for these light elements. C and N seem more abundant in close proximity to the holes (Figure S8A-B). A few instances of C and N within the “holes” are noted (Figure S8C-D), but it cannot be excluded that this spatial correspondence arises as an artifact due to noise. We segmented EDX maps and performed a Boolean intersection operation with the segmented “holes”. This showed that most “holes” are empty, although the presence of C and N in the segmented “holes” is considerably higher than that of Ca and P (Table S2). The lack of signal collected in most “holes” should be regarded as a technical limitation rather than true vacancy. In fact, as the lowest-contrast “holes” were the ones segmented and used in the Boolean intersection, this indicates that they may not contain enough matter to generate a strong EDX signal. If the “holes” were indeed empty pores, it would still be expected to collect C signal from the infiltrated embedding resin. Even if “holes” hosted non-collagenous organic substances, e.g., macromolecular complexes [19], C and N signals should still be detected. On the other hand, compositional information obtained from EELS experiments point towards the presence of C and N within the “holes” [42,43], corroborating their correspondence to collagen fibrils. Some additional technical considerations on EDX tomography in our work are provided in greater detail in Section 3.4.

### 3.3. Towards a better understanding of intra- and extra-fibrillar mineralization

#### Extra-fibrillar mineral is cross-fibrillar

Collagen fibrils are mostly in register across the entire volume in the tomograms where the banding pattern is visible in individual reconstructed slices (tomograms *i-a*/*b* and *ii-a*). Extra-fibrillar mineral, i.e., the phase not located within the gap zones, appears mostly aligned with collagen fibrils in tomograms *i-a*/*b* and *ii-a*, with mineral structures elongated along the direction of the collagen fibrils (Figure 6). We performed a “coarse” segmentation of the mineral phase based on grey-levels, followed by identification of disconnected sub-components. This showed that the segmented mineral regions are mostly interconnected, as in all tomograms the “Connected components – Multi-ROI” operation identifies a multi-ROI where a sub-ROI has significantly more voxels than the others (Table S3). This analysis also confirms the “cross-fibrillar mineralization” model proposed by Reznikov et al., where mineral aggregates form a continuous mineral phase extending beyond a single fibril, hence spanning adjacent fibrils [13]. The mineral phase encompassed in our segmentation appears to be mostly extra-fibrillar, since some intra-fibrillar mineral was excluded as the contrast in the corresponding gap zones was lower than our segmentation threshold (grey-level based). Therefore, in our reconstructions, the cross-fibrillar nature of the mineral can be accurately confirmed for the extra-fibrillar component.

**Figure 6.**
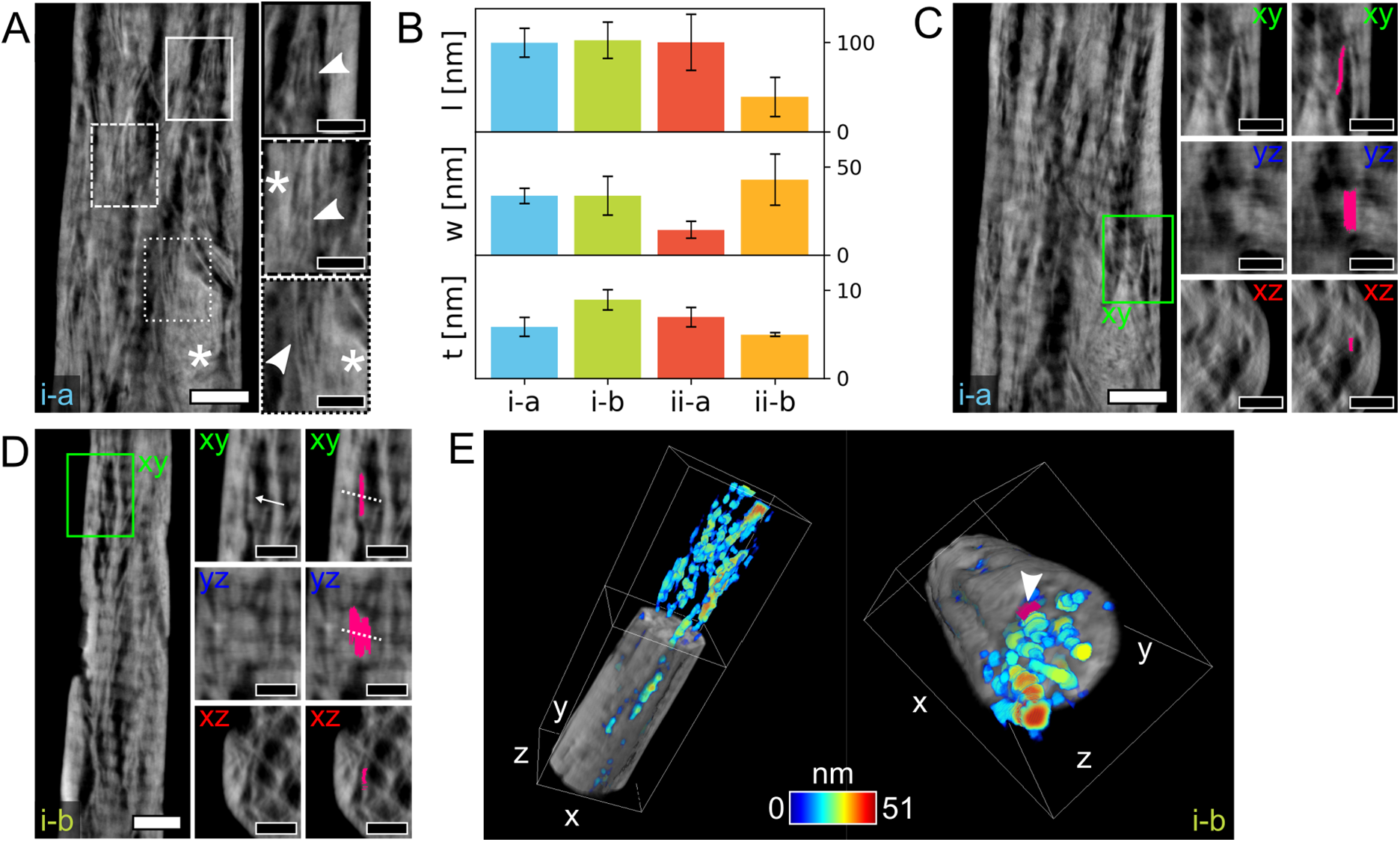
Representative examples of mineral plates. A) A representative reconstructed slice in the xy plane in tomogram *i-a* where both fine mineral structures (magnified in the insets, indicated by arrowheads) and larger aggregates (marked by *) are visible. B) Bar plots of average values of mineral length (l, top plot), width (w, middle plot), and thickness (t, bottom plot) for some representative mineral plates segmented in the four tomograms. C) An example of a mineral plate segmented in tomogram *i-a* (bright pink). Its plate-shape is confirmed by the cross-sectional view in the yz plane. In the xz plane, the mineral plate wraps around a “hole”. D) An example of mineral plate(s) segmented in tomogram *i-b* (bright pink). As the collagen banding pattern likely modulates the grey-levels in the entire slice, it is difficult to establish whether the segmented feature is a single plate or it is made of smaller plates fused together, especially when considering the cross-sectional view in yz (the potential boundary between sub-components is marked by the white arrow and dashed lines). In the xz plane, the mineral plate wraps around a “hole”. E) 3D renderings of tomogram *i-b* cropped to better show the segmented “holes” (heat map represents size). In the orientation on the right side, a mineral plate (same as in D) on the outside of a segmented “hole”, wrapping around it, can be noted (arrowhead). Scale bars are 200 nm in the overview images in A, C, and D, and 100 nm in their insets. A scale bar is not provided in E as the 3D representation is not an orthographic projection; the dimensions (x, y, z) of the white box are 555.55 × 1611.30 × 457.15 nm^3^.

#### Extra-fibrillar mineral is made of thin plates

In the complex scenario of fully mineralized tissues like mature bone, segmentation of individual mineral structures is not straightforward. In our reconstructions, some fine mineral structures were sometimes visible, but hard to isolate as they often seemed to fuse together (Figure 6A). In the extra-fibrillar mineral, a few of these structures appearing as one individual entity were manually segmented in each tomogram to offer some insights into their shape and size (Figure 6B, Table S4). Mineral structures in tomograms *i-a*/*b* and *ii-a* have a very similar length in the order of ∼100 nm. Mineral length appears smaller in tomogram *ii-b*, but this can be attributed to limitations in the segmentation of individual features given the high density of mineral throughout the tomogram, where small mineral structures appear extensively merged into bigger aggregates. The width of mineral plates is around 33-43 nm in tomograms *i-a*/*b* and *ii-b*, but smaller for tomogram *ii-a* (14 nm), while the plate thickness is in the 5-9 nm range. These values are in alignment with those found in the literature for 2D TEM measurements, typically in the order of 30-70 nm in length, 15-50 nm in width, and 5-10 nm in thickness [8,49–52], as well as for 3D measurements in early tomography studies (40-170 × 30-35 × 4-6 nm^3^) [53]. Additionally, the mineral we segmented appears to be extra-fibrillar, for which larger dimensions have been measured, with lengths up to 90-200 nm [6,8].

Over the years, varying size and shape have been reported for bone mineral. Discrepancies in mineral size can be mostly attributed to different techniques and/or sample preparation. For example, measurements from atomic force microscopy (AFM) yielded shorter crystals compared to TEM [54]. Mineral shape has also been often debated since early applications of TEM in bone research. While pioneering TEM analyses by Robinson showed the mineral to be plate-shaped (“tabular”) [49], others later proposed that the mineral is needle/rod-shaped [55]. This theory was disputed by others when imaging TEM samples at different tilt angles, concluding that the needle-shaped appearance was a misinterpretation arising from plates seen edge-on [56,57]. Using small-angle X-ray scattering, Fratzl et al. reconciled these views suggesting that mineral shape varies in different species, with human bone being more plate-shaped [58]. Later studies based on both TEM and AFM have consistently reported plate-shaped mineral [8,52,54,59], although the hypothesis of a needle-like habit has more recently resurfaced [13]. Our analyses overall support that the mineral is in the shape of thin plates, given the aspect ratio between the three dimensions (length/thickness, width/thickness, length/width). The plate morphology of the mineral is especially apparent when considering its shape in 3D and in distinct orthogonal planes (Figure 6C), further proving the advantages of 3D ET, especially on-axis tomography without artifacts, over 2D projection S/TEM images where edge-on views could lead to false interpretation of plates as needles.

#### Are extra-fibrillar plates made of even smaller components?

Given resolution limits and the small size of bone mineral, it is indeed possible that the segmented features correspond to aggregates of smaller sub-structures (e.g., platelets and/or needles), but their boundaries cannot be resolved, making them appear as one single feature. Similar challenges in identifying individual mineral structures have been reported in analyses of human trabecular bone [52]. Mineral structures (therein interpreted as needles) merging into larger aggregates were observed in human cortical bone, suggesting that bone mineral itself is a hierarchical assembly [13]. Early ET work also showed that mineral crystals fuse in a coplanar way to form large platelets [53], although it is worth noting that this study and several others dealing with collagen-mineral relationships in bone (where “bone” indicates a family of materials, and not just skeletal tissue) focused on the mineralized turkey tendon, which is a somewhat simplified system where collagen fibrils are well aligned with each other. Some authors also pointed out the existence of two populations of mineral structures in bone, i.e., small and large mineral components, with the large ones possibly being aggregates of small mineralites [54].

Interestingly, we noted some mineral plates presenting variation in grey-levels along their length (Figure 6D). This could indicate that they are composed of sub-structures, or, alternatively, this could be a modulation of the *Z*-contrast over the entire image due to the superimposition of the collagen banding pattern, as previously suggested [9,12]. For example, mineral plates splaying multiple bands, such as that shown in Figure 6D, were noted, but it remains difficult to establish whether they represent: i) the same purely extra-fibrillar entity (in agreement with [9,12]); ii) intra-fibrillar mineral growing in the inter-fibrillar space (as proposed in early ET studies by Landis et al. [53]); or iii) two distinct extra-fibrillar structures fused into one, in which case the distinction of different sub-components is irrelevant. Clearly, the direct visualization of bone mineral in fully mineralized bone is technically challenging, especially compared to simpler models like the mineralized turkey tendon. As bone mineral is the smallest biogenic crystal known, an even higher resolution is needed to clearly resolve mineral plates in ET, but this would inevitably reduce the volume analyzed, both in terms of field of view and sample thickness constraints.

#### Spatial relationship between extra-fibrillar mineral and “holes”

In tomograms *i-a*/*b*, some individual mineral plates appear surrounding the “holes” in the xz plane (Figure 6C-D). Given the evidence that “holes” are cross-sections of collagen fibrils, the presence of extra-fibrillar mineral wrapped around the fibril exterior yields a spatial relationship similar to the ultrastructural model described by Schwarcz [12]. This is even more noticeable in 3D, such as in Figure 6E, confirming the extra-fibrillar nature of the segmented mineral plate, which can be seen located on the outer surface of a segmented “hole” (i.e., collagen fibril).

#### Intra- and extra-fibrillar mineralization in EDX tomography

In reconstructed EDX maps, Ca and P are located in correspondence with both extra- and intra-fibrillar mineral (Figure 7A-B). These signals are more intense for the extra-fibrillar mineral, but they can also be detected within the gap zones, confirming intra-fibrillar mineralization (Figure 7C). This corroborates the contrast in the reconstructed HAADF signal, where extra-fibrillar mineral often appears brighter, while some gap zones display a lower contrast. Reconstructed EDX maps showing stronger Ca/P signal for the extra-fibrillar mineral are analogous to 2D STEM-EDX maps reported by McNally et al. [8]. While EDX tomography seems limited in the analysis of the organic phase, with low detection of C and N, it emerges as a viable tool to examine the mineral phase in 3D and reconstructed 2D slices.

**Figure 7.**
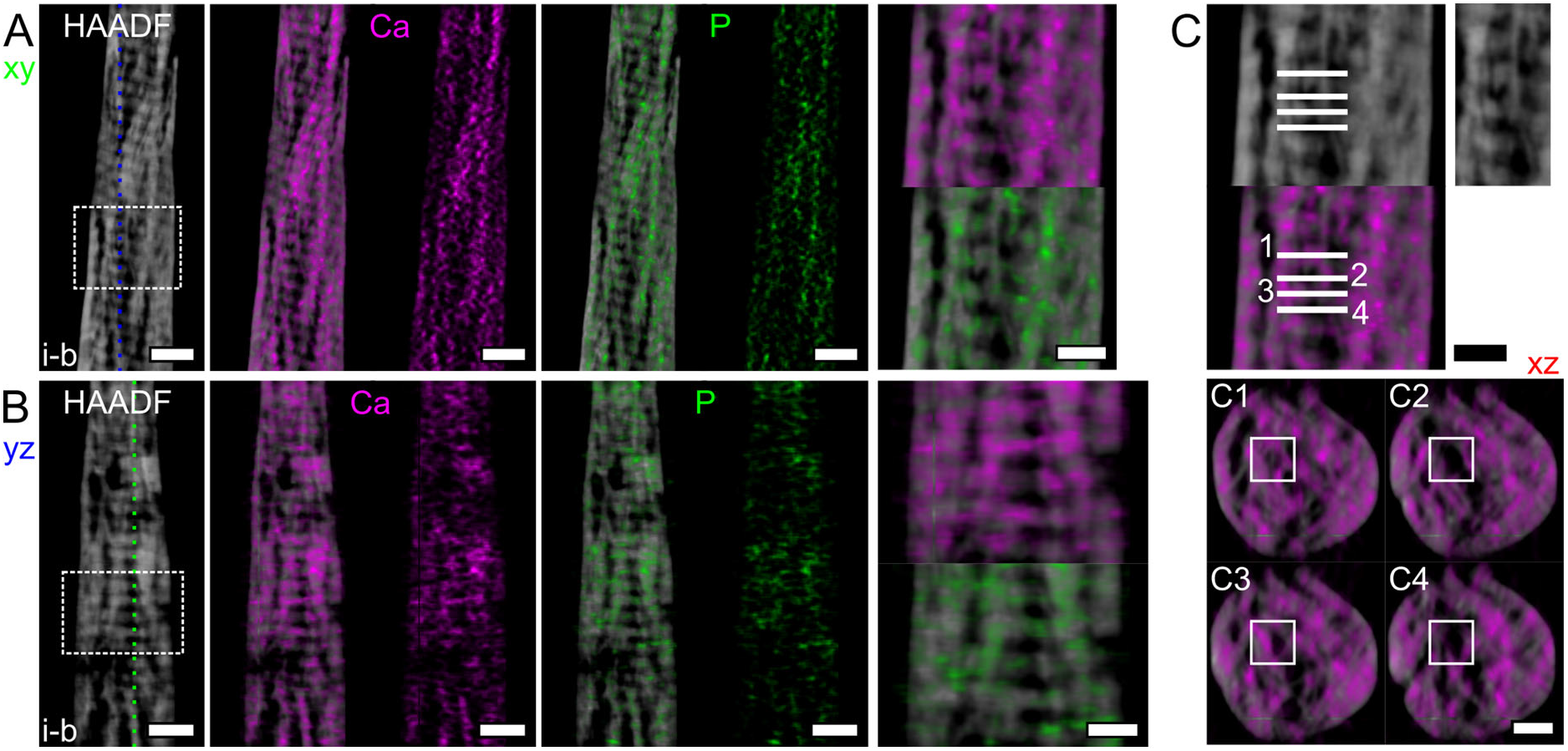
Representative examples of reconstructed HAADF-STEM slices and EDX maps. A) A representative reconstructed slice in the xy plane in tomogram *i-b* for the HAADF signal and EDX maps of Ca (magenta) and P (green) with and without the underlying HAADF slice. B) A representative reconstructed slice in the yz plane in tomogram *i-b* for the HAADF signal and EDX maps of Ca and P with and without the underlying HAADF slice. In both A and B, a magnified version of the area marked by the dashed rectangle is provided, where mineral can be seen both as extra- and intra-fibrillar. C) Different cross-sections (xz plane) are examined in gap (odd numbers) and overlap (even numbers) zones in the xy plane. From the reconstructed EDX maps of Ca in these different xz planes, intra-fibrillar mineralization, i.e., mineral in the gap zones, is evident. An unmarked version of the fibril examined is provided to the right of the HAADF slice in C, and marked by the rectangle in C1-4. Scale bars are 200 nm in A and B, and 100 nm in C and in the magnified areas corresponding to the dashed rectangles in B.

Not only does the intra-fibrillar mineral display a lower contrast (and Ca/P content) than the extra-fibrillar, but also the grey-level in each gap zone appears rather uniform. In other words, no individual mineral plates can be discerned in the intrafibrillar spaces, conversely to the extra-fibrillar mineral for which thin plates are observed. Some authors have proposed that the intra-fibrillar mineral is more amorphous [11,46], which would explain why individual crystals were not identified within the gap zones of our ET reconstructions. If intra- and extra-fibrillar mineral phases differ in crystalline structure, such a distinct property would challenge the inter-connection between these two phases.

### 3.4. Some technical considerations

HAADF-STEM ET has been applied to the study of bone ultrastructure [9,13,15,30] and various bone-biomaterial interfaces [28,29,60–62]. In sample *i-a*, we also acquired a tilt series in BF-STEM mode at the same time as the HAADF-STEM mode acquisition (Figure S9). Previous work has demonstrated both experimentally and by simulations that STEM tomograms of thick biological specimens (600 nm-1 μm) have a greater spatial resolution when acquired with a BF detector compared to a HAADF detector [63]. Our two rod-shaped samples (*i* and *ii*) have a diameter up to 700 nm, which is much thicker than conventional electron transparent wedge-shaped samples (100-200 nm). When comparing HAADF-STEM and BF-STEM tomography reconstructions, some features appear better resolved in the latter (Figure S9), suggesting that the BF detector may be more suitable for our thicker samples as reported in [63], even though differences are not striking. However, some artifacts (i.e., bright dots in the xy and yz planes and bright streaks in the xz plane) are present in the BF-STEM tomogram, deteriorating its overall quality (Figure S9). These artifacts were not present in the acquired nor aligned tilt series, but only in the individual reconstructed slices, implying that they originated during the SIRT reconstruction in the Inspect 3D software. It is possible that they are due to software computational assumptions related to the BF-STEM detector, or that the projection requirement is not entirely fulfilled, as incoherent image conditions for crystalline materials also depend on the BF detector semi-angle [63,64]. Therefore, in-depth analyses were performed only on HAADF-STEM tomograms.

EDX tomography is usually limited by a low signal-to-noise ratio (SNR), which requires long dwell times and/or high beam current to increase the dose, with the risk of inducing damage in the sample [65]. The S/TEM instrument used in this study (Talos 200X, Thermo Fisher Scientific) is equipped with a Super X-detector made of four separate silicon drift detectors, which allows for greater signal collection over a wider range of tilt angles compared to single detectors [40,66,67]. Additionally, the use of rotation tomography holders (in this work the Model 2050 by Fischione) eliminates X-ray collection issues related to the shadowing of the EDX detector [67]. Nonetheless, the spatial resolution in our EDX maps, especially for lighter elements like C and N, is not high enough to resolve the composition of some features of interest, such as the “holes”. Despite the not-so-optimal SNR, we kept a short acquisition time (∼2 min per map) to limit beam damage, which in fact was not visible during tilt series acquisition. It is also possible that the SNR was limited by X-ray absorption effects due to the large sample thickness. Based on our calculations using Eq. (1), up to nearly 40% X-rays could have been absorbed in the thicker parts of the sample (700 nm), especially for the lighter elements (C and N). More detailed simulations are needed to precisely quantify the fraction of X-rays absorbed and their path through the bone samples. In alternative to EDX, EELS tomography can be combined with HAADF-STEM tomography to correlate compositional and spatial information, as already demonstrated for bone-implant interfaces [29]. Greater spatial resolution is typically achieved in EELS than EDX [68], but as the signal collected in EELS is dependent on that transmitted through the sample, our thick samples were not suitable for this analysis.

Although on-axis ET requires specialized holders and rod-shaped samples, the advantages in the field of bone studies are not limited only to the removal of missing wedge artifacts. Using rod-shaped samples that were thicker than conventional wedge-shaped samples, we were able to collect more of the structural and compositional information contained in the sample thickness (z direction, considering xy as the image plane). This came at the cost of reduced information along the sample width (x direction), as the rod-shaped samples occupy only a limited portion of the field of view in the image plane (xy plane). However, the cylindrical, although slightly conical, shape of samples *i* and *ii* retains a near constant thickness at all tilts, unlike wedge-shaped samples, and its geometry better matches that of the features of interest present in bone at the nanoscale, i.e., the mineralized collagen fibrils. Collagen fibrils are reported to be oval-shaped in cross-section [44] and are routinely represented as rods [1,14]. At higher levels, fibrils organize themselves in bundles, which are described as cylindrical [1,14]. At the mesoscale, mineral aggregates are geometrically approximated in the shape of an ellipsoid [18,23,30], which is a solid of revolution.

## 4. CONCLUSIONS

Bone ultrastructure is challenging to assess with high resolution, elemental clarity, and over meaningful 3D volumes. In this work, we present a new method to characterize human bone ultrastructure and composition by correlating on-axis Z-contrast ET and EDX tomography in the TEM. Rod-shaped samples, with a diameter up to 700 nm, provided a larger sample thickness than conventional wedge-shaped TEM samples, as well as a geometry consistent with features of interest, such as mineralized collagen fibrils. Moreover, rotation of the rod-shaped samples on their long axis in on-axis ET removes missing wedge artifacts in the reconstructions. The segmentation of low-contrast spaces, the “holes”, debated to be some form of nano-porosity (eventually housing some non-collagenous organics) or collagen fibrils viewed in cross-section, revealed that they are rod-shaped with a diameter of ∼23 nm and are disconnected from each other. When examining mutually orthogonal planes in the reconstructions, collagen banding pattern is mostly seen in correspondence with the “holes”, supporting that these are cross-sectional views of collagen fibrils, especially along the overlap zones. 3D EDX confirmed both extra- and intra-fibrillar mineralization, but a clear understanding of their spatial relationship remains challenging. Extra-fibrillar mineral appears plate-shaped and richer in Ca and P than the intra-fibrillar mineral, with larger interconnected aggregates splaying multiple collagen fibrils. In this sense, the definition of an individual mineralized collagen fibril should perhaps be revisited, given the increasing evidence of extra-fibrillar mineral organized in a cross-fibrillar fashion.

## ACKNOWLEDGEMENTS

This study was supported by funds awarded to KG from the Natural Sciences and Engineering Research Council of Canada (NSERC) (grant no. RGPIN-2020-05722), the Ontario Ministry of Research, Innovation and Science (Early Researcher Award ER17-13-081), and the Canada Research Chairs Program from whom KG holds the Tier II Chair in Microscopy of Biomaterials and Biointerfaces. Scholarship support is acknowledged from the Ontario Graduate Scholarship and the Blanceflor Foundation for CM. Electron microscopy expe ents were performed at the Canadian Centre for Electron Microscopy (CCEM) at McMaster University (ON, Canada), a facility supported by NSERC and other government agencies. Funding support is also acknowledged from the Adlerbertska Foundation, the Kungliga Vetenskaps-och Vitterhets-Samhället i Göteborg, and the Svenska Sällskapet för Medicinsk Forskning (SSMF) for FAS, and from the Swedish Research Council (grant no. 2020-04715) and the Swedish state under the agreement between the Swedish government and the county councils, the ALF agreement (ALFGBG-725641), for AP. The Area of Advance Materials at Chalmers and at the Department of Biomaterials (University of Gothenburg) within the Strategic Research Area initiative launched by the Swedish government, and the Hjalmar Svensson Foundation are also acknowledged. The authors are thankful to Cheryl Quenneville for providing the bone sample, Dakota M. Binkley for the preparation of the bone sample, and Travis Casagrande for the preparation of the FIB samples.

## AUTHOR CONTRIBUTIONS

**Conceptualization:** All authors. **Formal analysis:** Chiara Micheletti. **Methodology**: Chiara Micheletti. **Investigation:** Chiara Micheletti. **Visualization:** Chiara Micheletti. **Supervision:** Furqan A. Shah, Anders Palmquist, Kathryn Grandfield. **Funding acquisition:** Kathryn Grandfield. **Writing – original draft:** Chiara Micheletti. **Writing – review and editing:** All authors.

## DATA AVAILABILITY STATEMENT

Data are provided in the main manuscript or in Supplementary Information.

