## Supplementary Information for "Shedding light (… electrons) on human bone ultrastructure with correlative on-axis electron tomography and energy dispersive X-ray spectroscopy tomography"

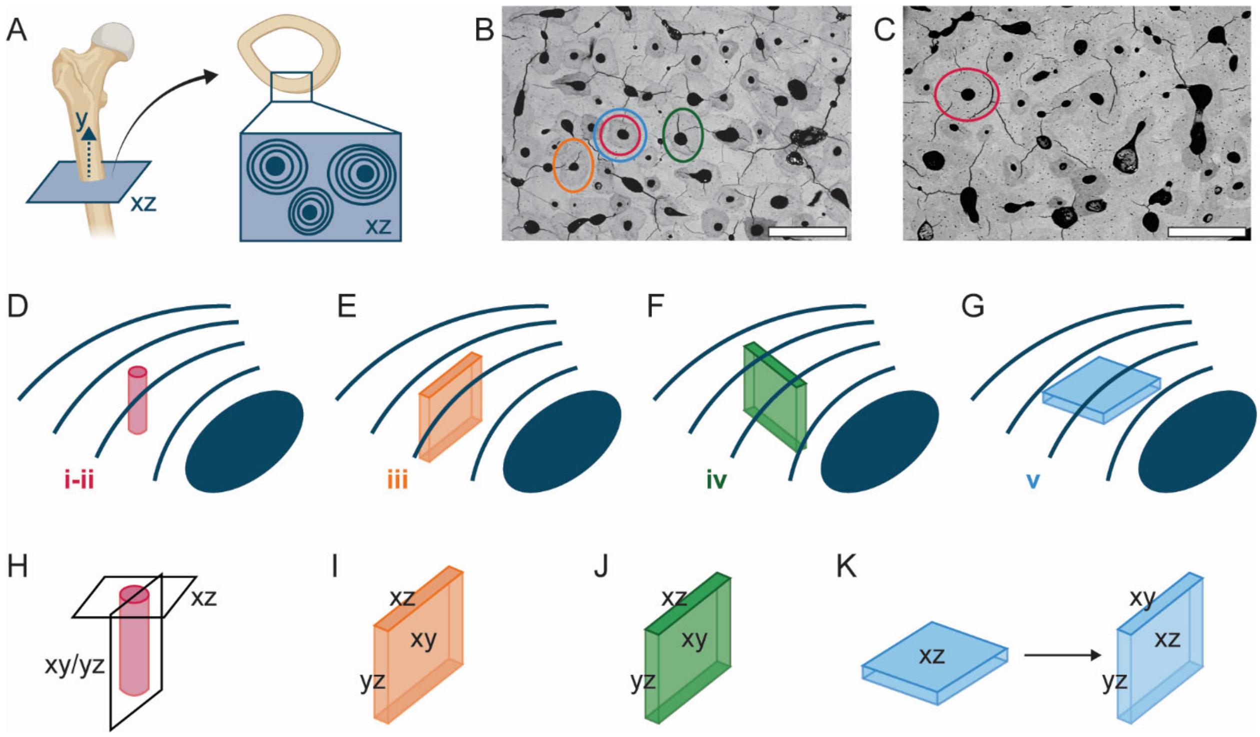


**Figure S1. Site selection and schematic representation of samples orientation.** A) Schematic representing the orientation of the "bulk" bone sample relative to the long axis of the femur. B-C) BSE-SEM images with circular marks around the osteons from which the samples were prepared. D-G) Orientation of both rod- (*i* and *ii* in D) and wedge-shaped (*iii* in E; *iv* in F*;* *v* in G) samples in relation to the osteonal axis and osteonal lamellae (the blue circle represents the Haversian canal, and the blue curved lines the inter-lamellar boundaries). H-K) Representation of the coordinate system convention used in the manuscript. The image plane in STEM corresponds to the xy plane, except for sample *v* where it corresponds to xz instead, as the sample was rotated during the *plan-view* FIB lift-out. Please note that given the cylindrical geometry of the rod-shaped samples, the definition of xy and yz planes is arbitrary, and depends on the insertion of the sample post in the holder and in the S/TEM instrument. In this case, the xy plane in the tomography reconstructions corresponds to the imaging plane during the tilt series acquisition. Scale bars are 500 µm in B and C. [Note: the drawings are not-to-scale, and the exact location of the samples from the Haversian canal is only indicative].

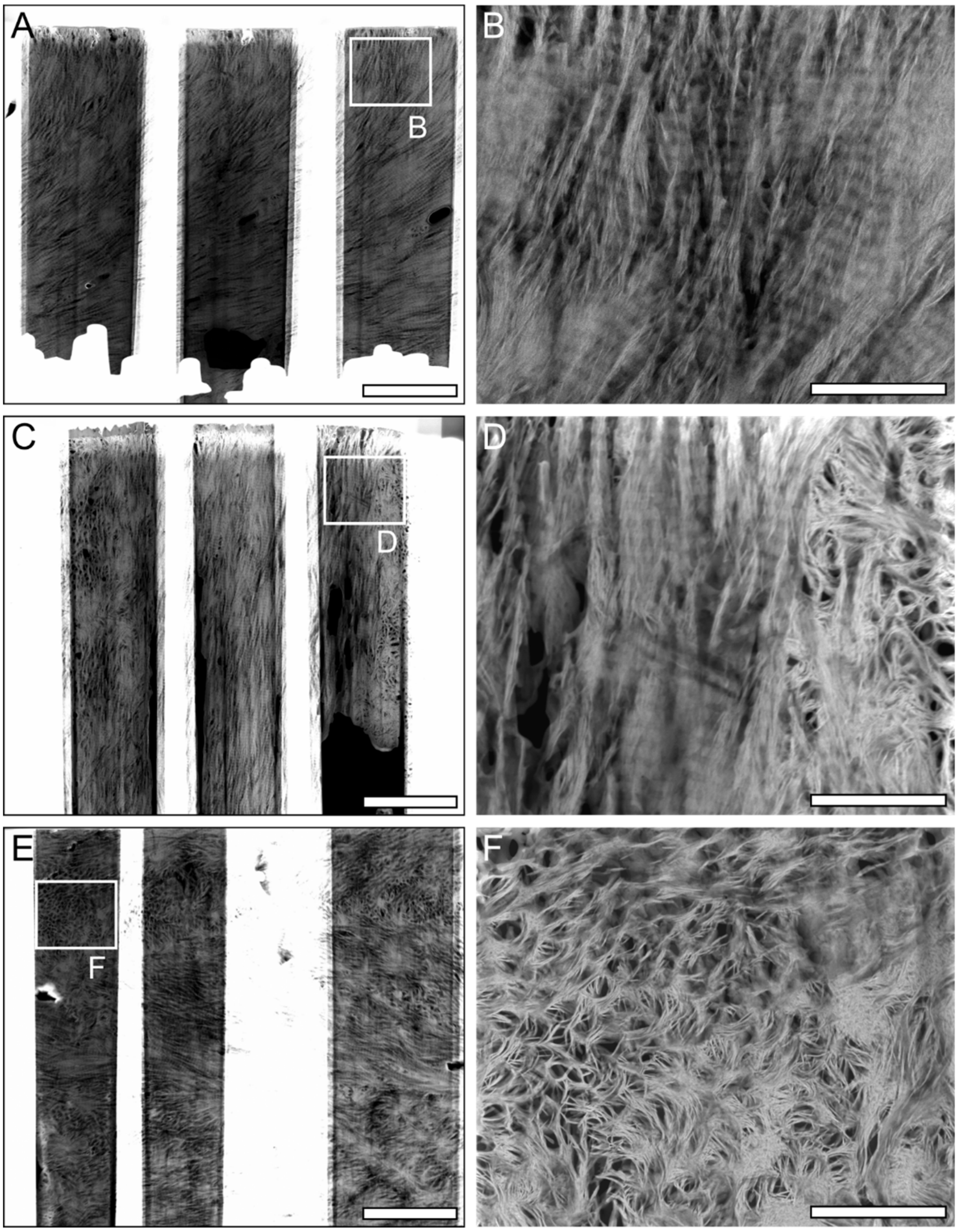


**Figure S2. HAADF-STEM images of wedge-shaped samples.** A) Overview image of wedge-shaped sample *iii*, where the characteristic banding pattern of in-plane collagen is predominant. B) Higher magnification view of the banding pattern. C) Overview image of wedge-shaped sample *iv*, where both longitudinal and lacy motifs are present. D) Higher magnification image showing the transition in orientation of collagen fibrils from in-plane (left side) to out-of-plane (right side). E) Overview image of wedge-shaped sample *v*, where collagen fibrils are mostly out-of-plane. F) Higher magnification image where “rosettes” and “holes” can be seen. Scale bars are 2 μm in A, C, and E, and 500 nm in B, D, and F. [Note: Figures S2A and S2F are also reported in the manuscript as Figures 2D and 2F, respectively].

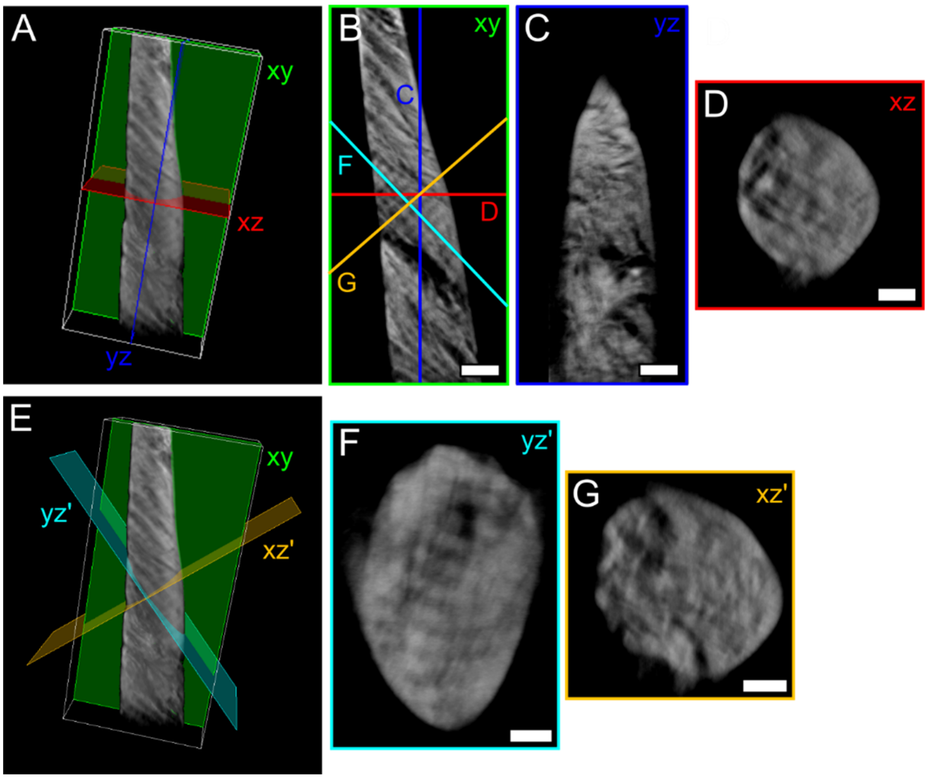


**Figure S3. Correspondence between longitudinal and lacy motifs.** A) 3D rendering of tomogram *ii-a*, showing three representative orthogonal planes, corresponding to xy (green), yz (blue), and xz (red), defined as per the coordinate system adopted in this work. B) A representative reconstructed slice in the xy plane where banding pattern is visible. C) Reconstructed slice in the yz plane corresponding to the blue line in A and B. Mineral-rich areas are present, but the banding pattern is not distinguishable. D) Reconstructed slice in the xz plane corresponding to the red line in A and B. E) 3D rendering of tomogram *ii-a*, showing three representative orthogonal planes, where yz' (cyan) and xz' (orange) are oriented perpendicular and parallel, respectively, to the banding pattern. F) Reconstructed slice corresponding to the cyan line in B and E, oriented perpendicular to the banding pattern, i.e., parallel to the collagen fibrils. The banding pattern is now clearly distinguishable, as opposed to C. G) Reconstructed slice corresponding to the orange line in B and E, oriented parallel to the banding pattern, i.e., perpendicular to the collagen fibrils. In both D and G, mineral-rich regions and "holes" can be observed. Scale bars are 200 nm in B, C, and F, and 100 nm in D and G. A scale bar is not provided in A and E as the 3D representation is not an orthographic projection; the dimensions (x, y, z) of the white box are 916.65 × 1943.55 × 579.60 $nm^3$ in both A and E.

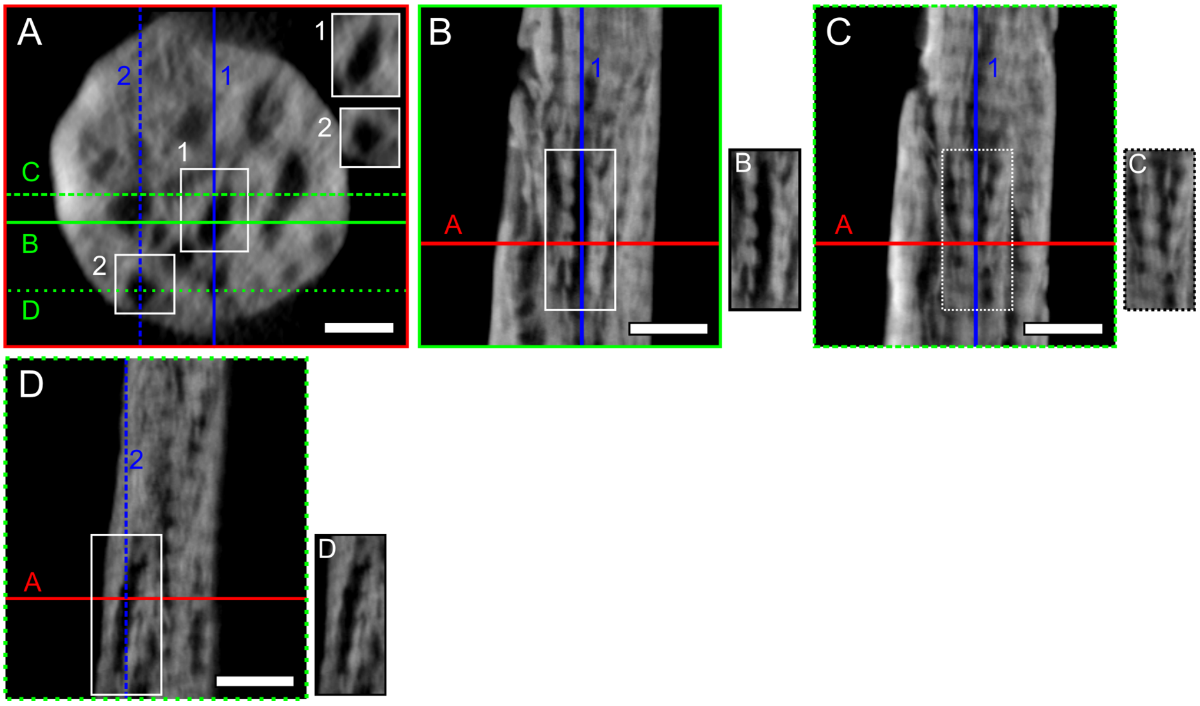


**Figure S4. Examples of "holes" where banding pattern is not apparent in cross-section.** A) A representative reconstructed slice in the xz plane in tomogram *i-b*. The "holes" marked by the white rectangles numbered "1" and "2" were examined in the xy planes shown in B/C and D, respectively. B) Reconstructed slice in the xy plane corresponding to the solid green line in A. The banding pattern is not clearly distinguishable within the "hole" (numbered "1" in A), but mostly around it. C) Reconstructed slice in the xy plane corresponding to the dashed green line in A. The banding pattern is more distinguishable than in B. The xy slices in B and C are 39.95 nm away from each other in the z direction. D) Reconstructed slice in the xy plane corresponding to the dotted green line in A. Collagen banding is barely visible, but the collagen fibrils could have been eroded by ion milling as this "hole" (numbered "2" in A) is approximately 13-15 nm away from the sample outer surface. An unmarked image of the regions marked by rectangles in B, C, and D is provided next to each panel. Scale bars are 100 nm in A, and 200 nm in B, C, and D.

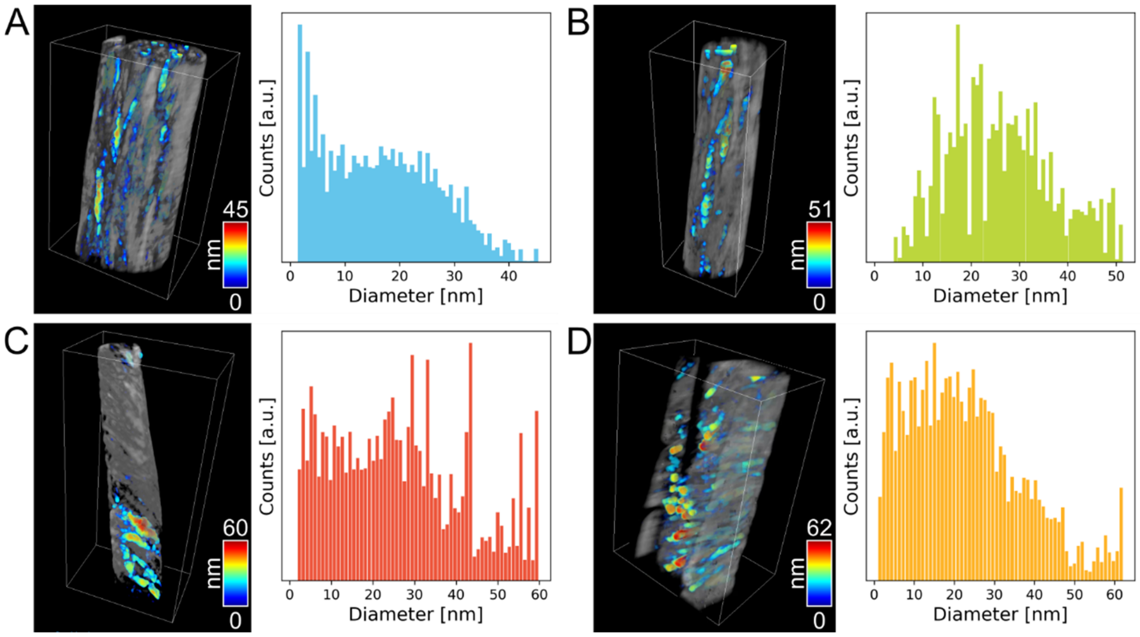


**Figure S5. 3D renderings of the tomograms with size analysis of the "holes".** The segmented "holes" are colour-coded based on size, and the distribution of the measured values is provided in the histogram: A) tomogram *i-a*; B) tomogram *i-b*; C) tomogram *ii-a*; D) tomogram *ii-b*. A scale bar is not provided as the 3D representation is not an orthographic projection; the dimensions (x, y, z) of the white box are 879.65 × 1416.20 × 620.50 $nm^3$ in A, 555.55 × 1611.30 × 457.15 $nm^3$ in B, 916.65 × 1943.55 × 579.60 $nm^3$ in C, and 1010.10 × 1515.52 × 704.48 $nm^3$ in D.

**Table S1.** Mean, median, minimum, and maximum "hole" size measured in each tomogram with "Volume thickness map" operation in the Dragonfly software.

| *Tomogram* | *Mean [nm]* | *Median [nm]* | *Minimum [nm]* | *Maximum [nm]* |
|---|---|---|---|---|
| *i-a* | 16.4 | 15.9 | 1.8 | 45.0 |
| *i-b* | 26.0 | 25.3 | 4.4 | 51.0 |
| *ii-a* | 26.4 | 24.5 | 2.3 | 59.3 |
| *ii-b* | 22.9 | 20.9 | 1.5 | 61.6 |
| ***Average*** | **22.9 ± 4.6** | **21.7 ± 4.3** | **2.5 ± 1.3** | **54.2 ± 7.7** |

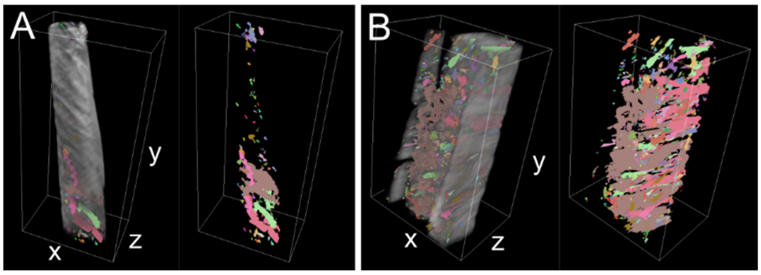


**Figure S6. Additional 3D renderings of the tomograms with segmentation of the "holes".** A) 3D rendering of tomogram *ii-a* with the segmented "holes", shown in different colours to represent their disconnected nature. B) Similar visualization as A for tomogram *ii-b*. A scale bar is not provided as the 3D representation is not an orthographic projection; the dimensions (x, y, z) of the white box are 916.65 × 1943.55 × 579.60 $nm^3$ in A, and 1010.10 × 1515.52 × 704.48 $nm^3$ in B.

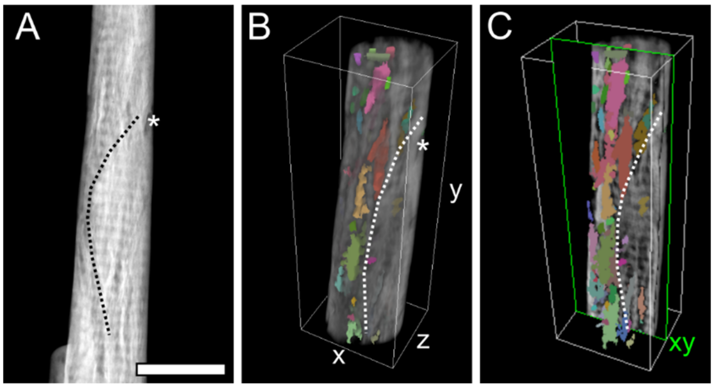


**Figure S7. “Holes” are more abundant around the periphery of a mineral ellipsoid.** A) HAADF-STEM image where the contour of a partial mineral ellipsoid (oriented longitudinally with respect to the long axis of the sample) is marked (black dotted line). B) 3D rendering of tomogram *i-b*, where the same mineral ellipsoid contour in A is marked by the white dotted line. For ease of comparison between A and B, a landmark is indicated by *. C) 3D rendering of tomogram *i-b* sliced along a representative xy plane where it becomes clearer that most segmented “holes” lie outside the mineral ellipsoid (contour marked by the white dotted line). Scale bar is 500 nm in A. A scale bar is not provided in B and C as the 3D representation is not an orthographic projection; the dimensions (x, y, z) of the white box are 555.55 × 1611.30 × 457.15 nm$^3$. [Note: Figures S7A and S7B are also reported in the manuscript as Figures 1C and 5C, respectively].

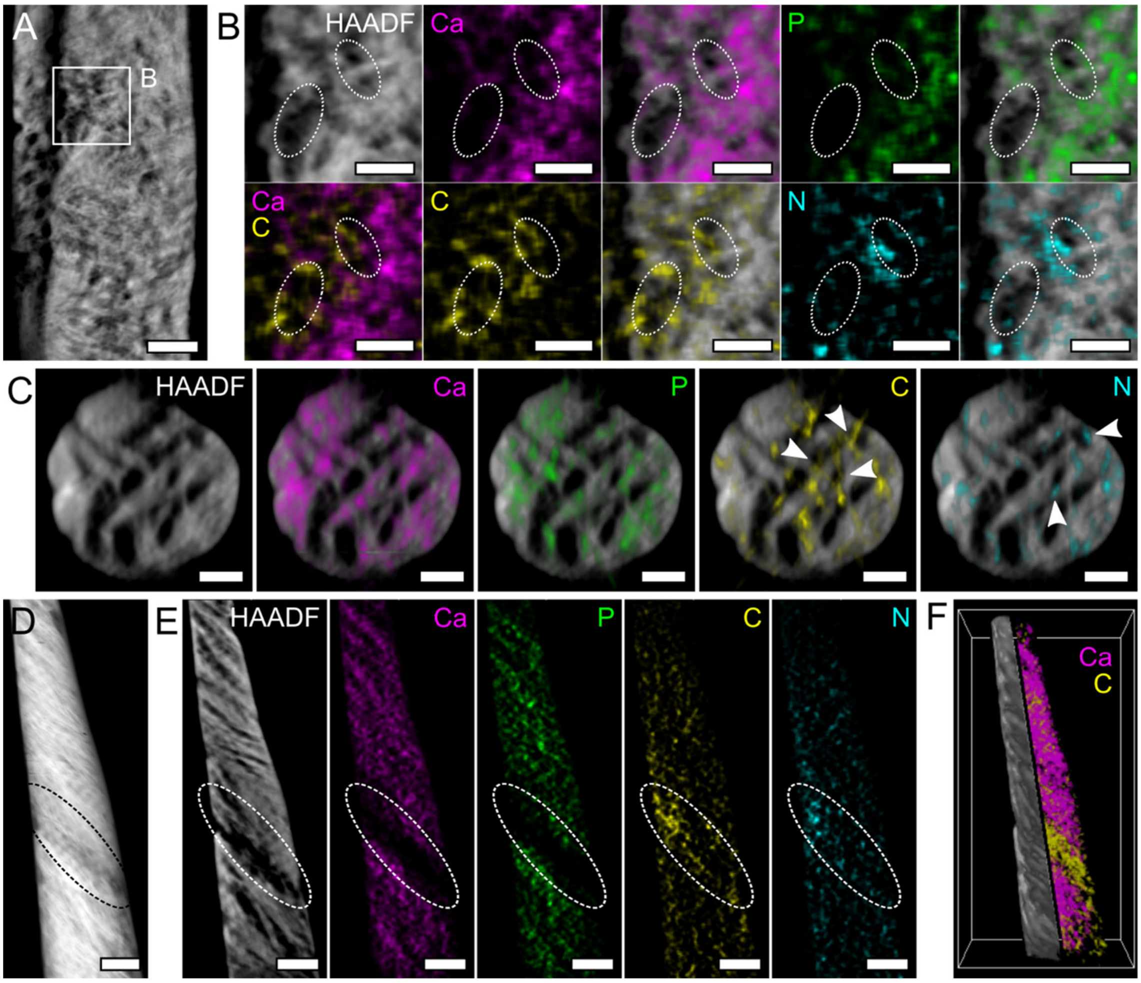


**Figure S8. Representative examples of reconstructed HAADF-STEM slices and EDX maps.** A) A representative reconstructed HAADF-STEM slice in the xy plane in tomogram *ii-b*. B) Reconstructed EDX maps (with and without underlying HAADF-STEM reconstructed slice) corresponding to the area marked by the white square in A. In the areas where "holes" are, Ca (magenta) and P (green) are mostly absent, while C (yellow) and N (cyan) signals appear more intense (e.g., regions circled by the white dotted lines). C) A representative reconstructed HAADF-STEM slice and EDX maps in the xz plane in tomogram *i-b*. While Ca and P are not detected within the "holes", C and N seem co-localized with some of them (arrowheads). D) HAADF-STEM image where tomogram *ii-a* was acquired. E) Reconstructed HAADF-STEM slice and EDX maps in the xy plane in tomogram *ii-a*. A region where faint banding pattern is present in the HAADF-STEM image in D and in the HAADF-STEM reconstructed slice appears depleted in Ca and P, but enriched in C and N. F) 3D rendering of HAADF-STEM and Ca/C signals on the left and right half, respectively. Scale bars are 200 nm in A, D, and E, and 100 nm in B and C. A scale bar is not provided in F as the 3D representation is not an orthographic projection; the dimensions (x, y, z) of the white box are 919.8 × 1869.0 × 579.6 $nm^3$. [Note: Figure S8D is also reported in the manuscript as Figure 1J, and it is cropped and duplicated here for ease of comparison with Figure S8E].

**Table S2.** Amount of Ca, P, C, and N contained within the segmented "holes", expressed as the number of voxels of each element with respect to the total number of voxels of the segmented "holes".

| *Tomogram* | *Ca [% voxels]* | *P [% voxels]* | *C [% voxels]* | *N [% voxels]* |
|---|---|---|---|---|
| *i-b* | 1.40 | 3.00 | 13.26 | 12.90 |
| *ii-a* | 0.02 | 0.12 | 10.08 | 16.37 |
| *ii-b* | 0.69 | 2.48 | 14.87 | 13.21 |

**Table S3.** Voxels labeled in the largest and second largest ROIs identified using the "Connected component – Multi-ROI" operation in the Dragonfly software applied to the segmentation of the mineral phase, and fraction of the largest ROI compared to the total number of voxels segmented for the entire mineral ROI.

| *Tomogram* | *Largest ROI [no. voxels]* | *Second largest ROI [no. voxels]* | *Fraction of largest ROI with respect to total ROI [%]* |
|---|---|---|---|
| *i-a* | 363,894,609 | 102,645 | 99.5 |
| *i-b* | 8,952,638 | 17,824 | 99.5 |
| *ii-a* | 87,083,428 | 94,695 | 99.4 |
| *ii-b* | 609,949,549 | 114,003 | 99.9 |

**Table S4.** Length, width and thickness of mineral plates segmented in each tomogram.

| *Tomogram* | *Length [nm]* | *Width [nm]* | *Thickness [nm]* |
|---|---|---|---|
| *i-a* | 81.9 | 40.2 | 5.3 |
| | 118.6 | 28.5 | 7.5 |
| | 92.6 | 31.4 | 5.2 |
| | 90.6 | 33.6 | 6.3 |
| | 115.2 | 34.3 | 4.9 |
| ***Average*** | **99.8 ± 16.2** | **33.6 ± 4.3** | **5.8 ± 1.1** |
| *i-b* | 86.1 | 28.7 | 8.9 |
| | 96.5 | 41 | 10.5 |
| | 100.9 | 24.6 | 7.6 |
| | 137.1 | 24.6 | 8.1 |
| | 90.7 | 49.2 | 9.6 |
| ***Average*** | **102.3 ± 20.3** | **33.6 ± 11.0** | **8.9 ± 1.1** |
| *ii-a* | 111.7 | 11.6 | 5.7 |
| | 64.4 | 20.0 | 7.3 |
| | 123.8 | 11.6 | 7.9 |
| ***Average*** | **100.0 ± 31.4** | **14.4 ± 4.8** | **7.0 ± 1.1** |
| *ii-b* | 64.4 | 53.5 | 5.1 |
| | 27.4 | 48.8 | 5.1 |
| | 25.2 | 26.3 | 4.7 |
| ***Average*** | **39.0 ± 22.0** | **42.9 ± 14.5** | **4.7 ± 0.2** |

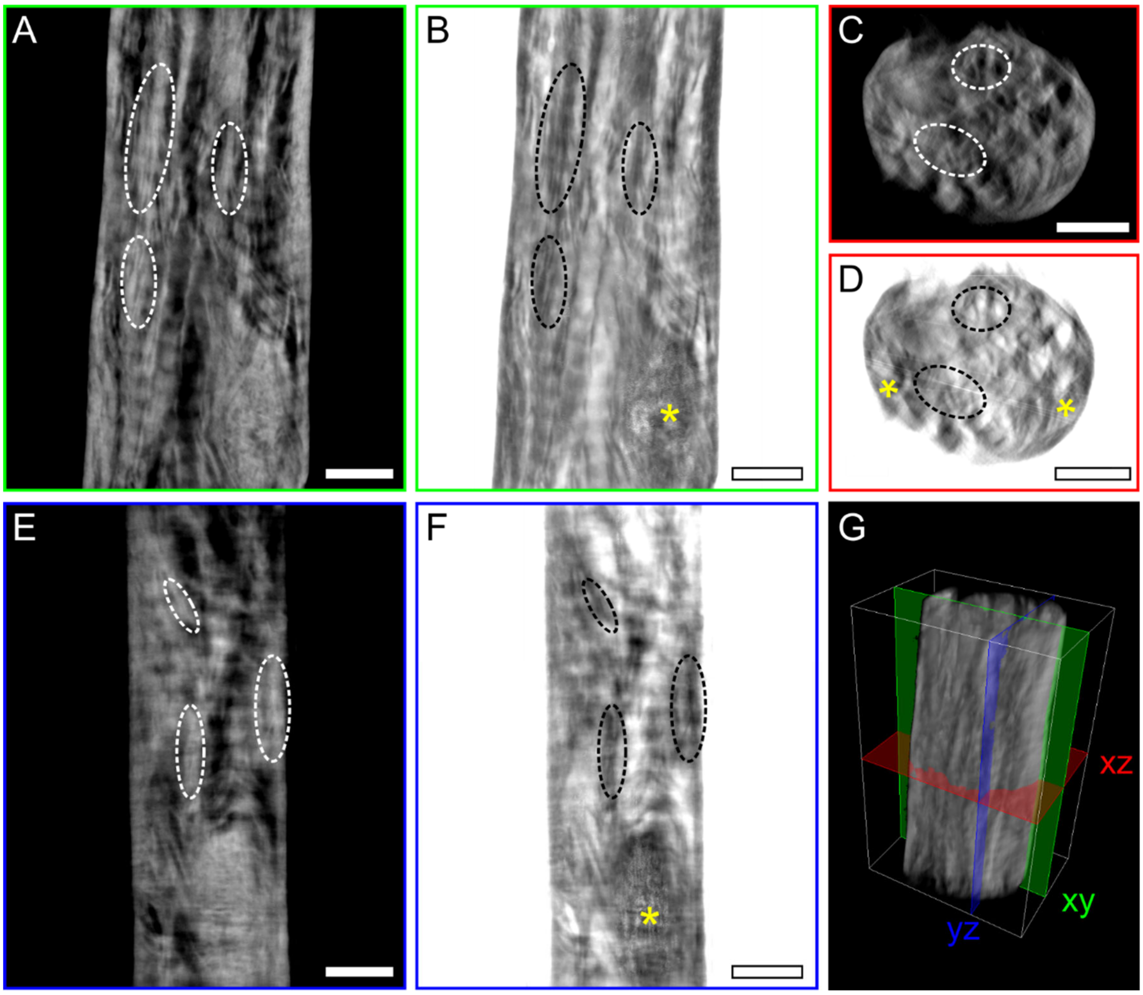


**Figure S9. Comparison between reconstructed slices acquired in HAADF-STEM *vs.* BF-STEM mode.** A-B) A representative reconstructed slice in the xy plane (green) acquired in HAADF-STEM (A) and BF-STEM (B) mode. C-D) A representative reconstructed slice in the xz plane (red) acquired in HAADF-STEM (C) and BF-STEM (D) mode. E-F) A representative reconstructed slice in the yz plane (blue) acquired in HAADF-STEM (E) and BF-STEM (F) mode. Although differences are not striking, mineral structures appear overall less blurred and better distinguishable from the background in all planes in BF-STEM mode. Some examples are marked by the dotted ovals. Artifacts are present in the BF-STEM reconstruction in the form of bright dots in the xy and yz planes and bright streaks in the xz plane (regions marked by the yellow *). G) 3D rendering of the HAADF-STEM tomogram showing the position of the three representative slices in A-F. Scale bars are 200 nm in A-F. A scale bar is not provided in G as the 3D representation is not an orthographic projection; the dimensions (x, y, z) of the white box are 879.65 × 1416.2 × 620.5 $nm^3$.
